# The circadian molecular clock in the suprachiasmatic nucleus is necessary but not sufficient for fear entrainment in the mouse

**DOI:** 10.1101/2023.06.26.546624

**Authors:** Ivana L. Bussi, Miriam Ben-Hamo, Luis E. Salazar Leon, Leandro P. Casiraghi, Victor Y. Zhang, Alexandra F. Neitz, Jeffrey Lee, Joseph S. Takahashi, Jeansok J. Kim, Horacio O. de la Iglesia

## Abstract

Nocturnal aversive stimuli presented to mice during eating and drinking outside of their safe nest can entrain circadian behaviors, leading to a shift toward daytime activity. We show that the canonical molecular circadian clock is necessary for fear entrainment and that an intact molecular clockwork in the suprachiasmatic nucleus (SCN), the site of the central circadian pacemaker, is necessary but not sufficient to sustain fear entrainment of circadian rhythms. Our results demonstrate that entrainment of a circadian clock by cyclic fearful stimuli can lead to severely mistimed circadian behavior that persists even after the aversive stimulus is removed. Together, our results support the interpretation that circadian and sleep symptoms associated with fear and anxiety disorders may represent the output of a fear-entrained clock.

**One-Sentence Summary:** Cyclic fearful stimuli can entrain circadian rhythms in mice, and the molecular clock within the central circadian pacemaker is necessary but not sufficient for fear-entrainment.

## Introduction

The coding of threatening and aversive stimuli as fear represents a highly conserved adaptation shared across most animals, including humans. For prey species, the ability to encode complex spatial and temporal predator cues, whether innate or learned, is an essential survival mechanism. Likely, the 24-h structure of a predator’s activity serves as a crucial temporal cue to its prey. Studies on wild animal populations have indeed shown that the 24-h activity patterns of prey species can be influenced by their predators’ activity patterns (*1-6*). While the mechanisms underlying this temporal predator avoidance are unknown, we have previously shown that nocturnal fear can entrain circadian rhythms of foraging and feeding. Rats living in a safe nest, when required to obtain food and water by venturing into a separate foraging area, predominantly feed and drink during their usual nocturnal phase. However, when the foraging area is made dangerous through randomly distributed footshocks during the dark phase of the light-dark (LD) cycle, the rats shift their feeding and drinking activities to the daytime (*7*). This unusual diurnal behavior observed in a nocturnal rodent could have emerged either as an avoidance response to nocturnal footshocks or as a result of learning wherein the light phase becomes conditionally associated with safety. Alternatively, our studies showed that rats can anticipate the safe light phase and begin venturing into the foraging area, initiating eating and drinking activities at the end of the dark phase, before lights turn on signaling the safe time of day. This finding suggested the potential involvement of a circadian clock in predicting the cyclic aversive stimulus. Indeed, we have shown that rhythms of activity in the foraging area and feeding resulted from the entrainment of a circadian oscillator (*7*). The reliance of this oscillator on the canonical molecular circadian clock and the timing by the central circadian pacemaker located within the suprachiasmatic nucleus (SCN) is yet to be determined. Here, we exploit our fear-entrainment paradigm in mice to (a) show that cyclic fear entrainment is likely a conserved feature of the circadian system of mammals, (b) reveal basic properties of entrainment by cyclic fear, (c) confirm that the fear-entrained oscillator depends on the canonical molecular clock, and (d) demonstrate that an intact molecular clock within the SCN is necessary, though not sufficient, for fear entrainment. Our findings underscore the salience of cyclic 24-h fear stimuli as a key entraining environmental cycle for the circadian system that has the ability to drastically shift the temporal distribution of behavior, providing a new neural framework to understand circadian and sleep disruptions associated with fear and anxiety disorders.

## Results

### Nocturnal fear entrains circadian rhythms of foraging and feeding behavior in the mouse

To assess whether a fearful stimulus can entrain circadian rhythms of locomotor activity in the mouse, we first built customized cages that mimic a more naturalistic environment than regular housing cages. These cages have two different compartments: a nesting compartment where the mice are safe from the aversive stimulus and a foraging area where food and water are available ad libitum but where footshocks can be delivered through a metal grid floor (Fig. S1A). These cages allowed us to record three different behavioral outputs: locomotor activity within the nesting area (hereafter referred to as *nest activity*) and within the foraging area (hereafter referred to as *foraging*) with IR detectors, and feeding through a nose-poke detector in the feeder. Adult male and female mice were housed in these customized cages and subjected to a 12:12 light-dark (LD) schedule. After 10 days in baseline conditions, we started delivering three footshocks per hour—randomly distributed throughout time—in a 12-h time window within either the dark (dark fear = DF) or the light (light fear = LF) phase of the LD cycle. LF mice displayed a mild change in behavior after the presentation of the shocks, avoiding the daytime activity evident during baseline (Fig 1A left panel, Fig. S1B). In contrast, DF mice shifted their nocturnal activity to the light phase, with most of the activity occurring during the first hours of the light phase (Fig. 1A right panel, Fig. S1E). DF mice also showed increased activity during the last hour of darkness before lights-on, which we interpret as anticipation of the shock-free phase. Waveform analysis including all animals in each group confirmed the nocturnal and diurnal activity patterns, respectively, of LF and DF animals (Fig. 1B, Fig. S1C, F). To test whether the daytime activity of DF mice represented an acute response to the dark-phase shocks, we released DF mice into constant darkness conditions (DD) in the absence of footshocks (hereafter referred to as post-shocks). Upon release into constant conditions, the phase of the rhythms of foraging, feeding, and, to some extent, nest activity resembles the phase during the presentation of the shocks, indicating that the daytime activity was the result of authentic entrainment by the nocturnal footshocks (Figs. 1A right panel, Fig. S1E). Waveform analysis of all three behaviors confirmed the phase of DF mice after their release into constant conditions (Fig. 1B right panel, Fig. S1F).

**Figure 1.**
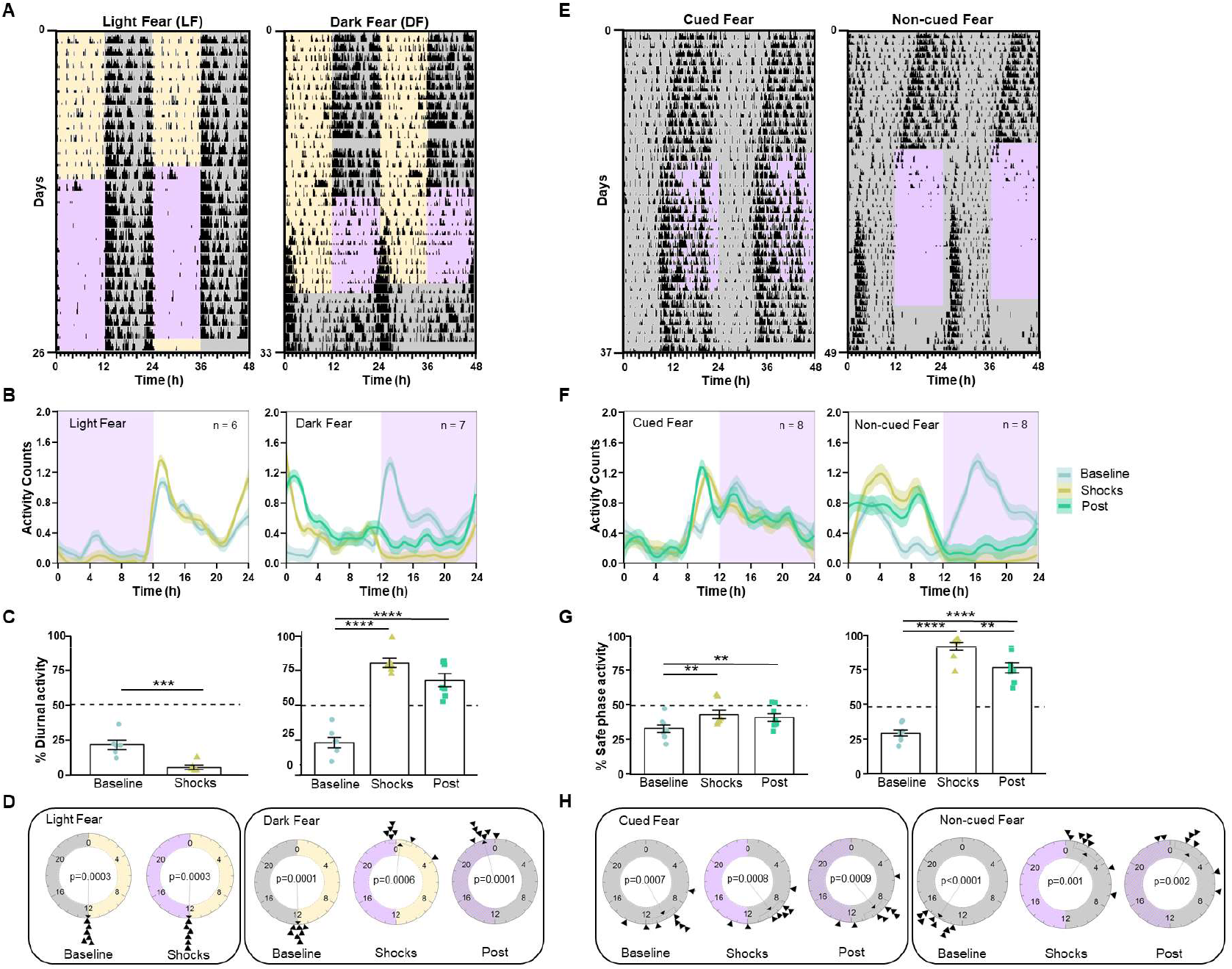
Cyclic fear entrains a circadian oscillator under a light-dark cycle or constant darkness. **(A)** Representative foraging actograms from mice in LD subjected to cyclic fear presented either during the day (LF, left) or during the night (DF, right). Yellow and gray shading, respectively, represents the light and dark phases of a 12:12 LD cycle. Purple shading represents the 12-h window of time at which 3 footshocks/hour randomly distributed over time were presented. The DF mouse is released into constant conditions (DD and no shocks). **(B)** Average foraging activity patterns (line represents locally estimated scatterplot smoothing (LOESS) regressions obtained from the mean values of all mice mean and shading SEM) from mice under LF (left, n=6) or DF (right, n=7). **(C)** Percent of activity that took place during the daytime or extrapolated daytime across the different experimental stages from the same mice shown in **B**. Bars represent the mean ± SEM. **(D)** Rayleigh plots representing the time of activity onset across the different experimental stages for mice subjected to LF (left) or DF (right). **(E)** Representative foraging actograms from mice in DD subjected to either cued (left) or non-cued (right) fear during the subjective night. Purple shading represents the shock-delivering window **(F)** Average foraging activity patterns from mice subjected to cued (left, n=8) or non-cued fear (right, n=8). **(G)** Percent of activity that took place during the safe phase (window of time without shocks) or extrapolated safe phase across the different experimental stages from the same mice shown in **F. (H)** Rayleigh plots representing the time of the activity onset during the different stages of the protocol for mice subjected to cued (left) or non-cued fear (right). Asterisks indicate statistically significant differences according to Tukey comparisons following LMM analysis (Table S1): ** p < 0.01, *** p < 0.001.

We analyzed these data as follows: First, we determined the percent of diurnal activity for each of the three behavioral outputs within each animal. Then, we analyzed this variable through linear models with mixed effects (LMM), with the group (LF vs. DF) and experimental stage (baseline, shocks, or post-shocks) as fixed factors and the individual mouse as a random factor. (Table S1). This analysis revealed that LF mice displayed less diurnal activity during the presentation of shocks than during baseline. For instance, 21.4 ± 3.3% (mean±SEM) of the foraging activity occurred during the daytime on baseline, but only 5.4± 1.5% occurred during the daytime when shocks were delivered. The analysis also revealed that DF animals switched from being predominantly nocturnal in all three behaviors during baseline to being predominantly diurnal during the presentation of shocks as well as upon their release into constant conditions. For instance, 23.3 ± 3.7% of the foraging activity occurred during the daytime on baseline, 80.5 ± 4.7% occurred during the daytime when shocks were delivered, and 68.1 ± 4.7% during the projected daytime after their release into constant conditions. Second, we used circular statistics to assess the changes in the phase of the 24-h onset of foraging activity. Circular plots of the onset of each animal’s foraging time are presented in Fig. 1D, and their analysis through the Rayleigh test is in Table S2. This analysis revealed that LF animals started foraging at the time of lights-off during baseline, and this phase remained during the presentation of shocks. In contrast, although DF animals also started foraging at the time of lights-off during baseline, they shifted by approximately 12 hours during the presentation of shocks, with the phase of the foraging start time occurring around lights-on and coincident with the termination of the daily shocks. Importantly, this latter phase remained when the animals were released into constant conditions.

### Cyclic fear entrains circadian rhythms of foraging and feeding behavior under constant darkness conditions

Our fear-entrainment paradigm under LD conditions presents the confounding effect that footshocks in both groups are always paired with a particular phase of the LD cycle, raising the possibility that cyclic fear could only entrain circadian rhythms when an external time reference is present. To test whether this is the case, we repeated our experiment under DD conditions. During a ∼12-day baseline phase in which animals were placed in the fear chambers under DD, mice displayed the typical <24-h circadian period in rhythms of foraging, feeding, and nest activity (Fig. 1E, Fig. S2A, D). After the baseline phase, the control group received three footshocks per hour—randomly distributed throughout time—during a 12-h time window, but these shocks were preceded by a 20-sec tone (*cued-fear*), which allowed the animals to predict the arrival of the footshock. The experimental group received the same temporal distribution of footshocks, but the shocks were not paired with a tone (*non-cued-fear*). The results showed that cued-fear mice effectively predicted and avoided the shocks and did not change the circadian phase of any of the three behavioral outputs measured (Fig. 1E left panel, Fig. S2A). In contrast, non-cued-fear animals shifted their rhythms within a few cycles, leading to a pattern of foraging, feeding, and nest activity that effectively avoided activity during the shock phase (Fig. 1E right panel, Fig. S2D). Importantly, upon release into constant conditions (with no shocks), both groups displayed a circadian phase that was predicted by the phase displayed during the presentation of shocks. Waveform analysis confirmed the 24-h temporal pattern of all three behaviors in both cued and non-cued mice (Fig. 1F, Fig. S2B, E).

We determined the percent of activity for each of the three behavioral outputs that took place during the safe phase (the 12-h window without footshocks) within each animal and analyzed the change in this variable across stages through an LMM. In cued-fear animals, during the presentation of shocks, the percent of each behavior displayed during the safe phase differed slightly relative to the percent activity in the extrapolated phase during baseline, and continued to differ slightly from baseline when the shocks were removed (Fig. 1G left panel, Fig. S2C, Table S1). These slight changes in phase were likely the consequence of the fact that the animals exhibited a different period than the 24-h period of the cyclic cued shocks. In contrast, the percent of activity during the safe phase in mice subjected to the non-cued fear protocol changed dramatically during the presentation of shocks (94.6% for foraging, 96.4% for feeding, and 63.5 for nest activity) when compared to the percent of activity in the extrapolated phase during the baseline (30.5% for foraging, 30.9% for food, and 36.2% for nest activity). These results clearly revealed that all three behaviors were largely restricted to the safe phase of the 24-h shock cycle. The trend persisted when shocks were removed, with a large percent of each behavior restricted to the extrapolated safe phase during the post-shocks stage (78.7% for motion, 84.9% for food, and 50.5 for nest activity; Fig. 1G right panel, Fig. S2F, Table S1). Circular plots followed by the Rayleigh tests clearly showed that while the phase of rhythmic foraging in cued-fear animals did not shift with the presentation of shocks, the rhythm was readily entrained by the presentation of shocks in non-cued fear animals, in which the onset of activity occurred right after the time the shock period ended (Fig. 1H, Table S2).

### Circadian clock gene expression in the suprachiasmatic nucleus is not entrained by cyclic fear

Entrainment of circadian rhythms by cyclic nocturnal fear could be the result of entrainment of the central circadian clock housed within the SCN. To test this possibility, we performed *in situ* hybridization for the clock genes *Bmal1* and *Per1* in coronal brain slices from either LF or DF mice euthanized every four hours throughout the 24-h cycle. The pattern of *Bmal1* and *Per1* expression in both groups was indistinguishable and showed the expected circadian expression that would result from photic entrainment of the SCN, i.e., respectively high and low expression of *Per1* and *Bmal1* during the light phase (Fig. 2). Cosinor analysis followed by a Wald test confirmed there were no detectable differences in the amplitude or phase of clock gene expression patterns between LF and DF animals (Table S3). Because of their involvement in contextual fear-conditioning, we also examined the pattern of clock gene expression in the dentate gyrus of the hippocampus (DG) and basolateral amygdala (BLA). The cosinor analysis failed to detect any rhythm in the expression of the *Per1* gene in the BLA or DG for LF or DF samples (Fig. S3, Table S3).

**Figure 2.**
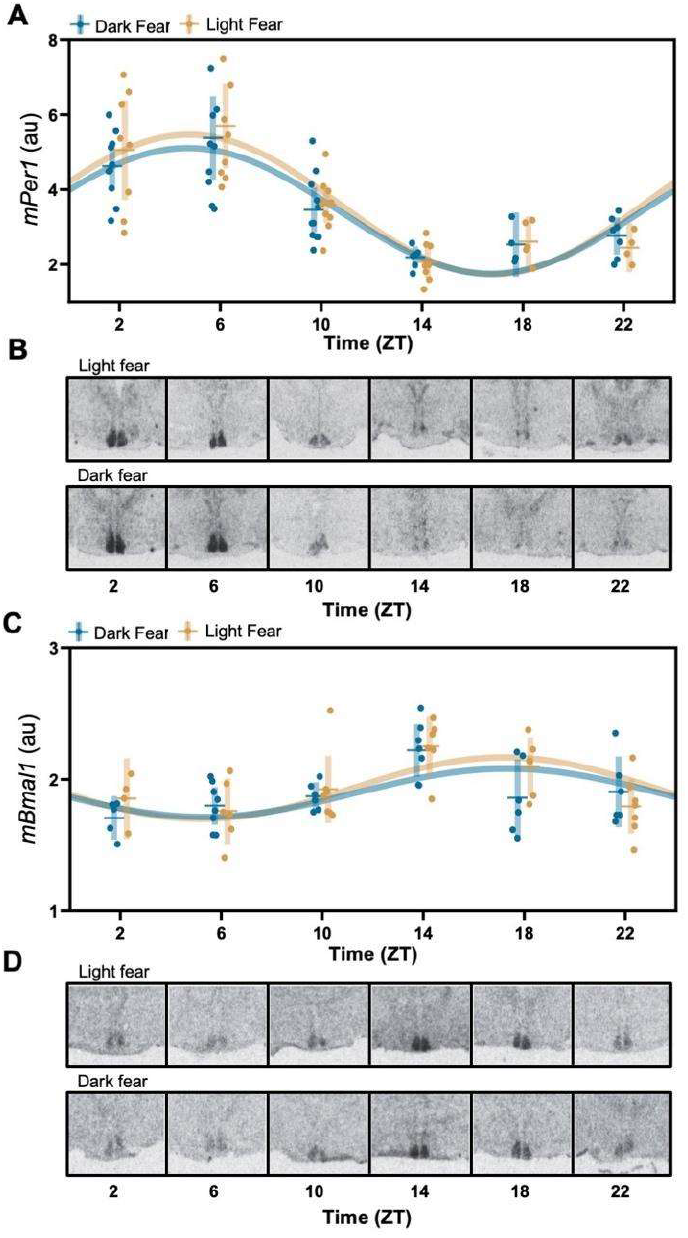
Circadian clock gene expression in the SCN is entrained to the light-dark cycle independently of fear entrainment. **(A, C)** Daily patterns of *mPer1* and *mBmal1* mRNA expression, respectively, in mice housed under a 12:12 LD cycle subjected to LF or DF. Each dot represents an individual mouse, horizontal and vertical lines, respectively, represent the mean and SEM. The best-fitting sine wave with a 24-h period for each group is presented for illustrative purposes (solid lines, fit parameters are presented in Table S3). **(B, D)** Representative autoradiographs of coronal brain sections at the level of the anterior hypothalamus, hybridized with a radioactive probe for *mPer1* and *mBmal1* mRNA detection, respectively.

### The expression of the clock gene *Bmal1* in the forebrain is necessary for fear entrainment

The master regulation of overt behavioral and physiological rhythms in mammals relies on the transcription-translation feedback loop of canonical clock genes within cells of the SCN (*8, 9*). However, several studies have shown that behavioral circadian rhythms entrained to time-restricted food access do not rely either on the SCN or the canonical molecular clockwork (*10-12*). We reasoned that this independence from the SCN molecular clockwork could be a characteristic of non-photically entrained circadian rhythms and examined whether the circadian canonical clock is necessary to sustain fear-entrained circadian rhythms. We tested mice lacking the clock gene *Bmal1* in the forebrain, which readily entrain to time-restricted food access (*12*), in our fear-entrainment paradigm. In this mouse line, a Cre-driver under the promoter of the *CamKII* gene targets the deletion of *Bmal1* to neurons within structures within the forebrain and leads to rhythmic behavior under LD driven by masking—the expression of rhythmic behavior as mere response to the LD cycle—, but to arrhythmic locomotor activity patterns under DD conditions in mice lacking both copies of the gene. However, a single copy of *Bmal1* in the forebrain is sufficient to sustain behavioral circadian rhythmicity (*12*).

We tested mice lacking either both copies of *Bmal1* (*Cami-Bmal1*^-/-^), only one copy of *Bmal1* (*Cami-Bmal1*^*+/-*^) or none (*Cami-Bmal1*^*+/+*^) under DF and non-cued fear protocols. In LD and during baseline conditions, *Cami-Bmal1*^-/-^, *Cami-Bmal1*^*+/-*^ and *Cami-Bmal1*^*+/+*^ displayed nocturnal activity. When subjected to DF, the three groups successfully shifted their behaviors to the light phase, indicating that the lack of *Bmal1* expression does not impair fear perception. Upon release into constant conditions (DD and no footshocks), *Cami-Bmal1*^*+/-*^ and *Cami-Bmal1*^*+/+*^ mice maintained the phase acquired during the shock-presentation phase (Fig. 3A, Fig. S4A). In contrast, *Cami-Bmal1*^-/-^ displayed an arrhythmic pattern of behavior, suggesting that the synchronization during the shock presentation phase was the result of pairing the aversive stimulus to the dark phase instead of the entrainment of a functional circadian clock (Fig. 3A and Fig. S4A, left and center panels). Waveform analysis confirmed the results for each group, and also revealed that the *Cami-Bmal1*^-/-^ group displayed lower levels of foraging during the presentation of shocks than after release into constant conditions (Fig. 3B). The LMM analysis of the percent of each behavior that took place during the daytime or the extrapolated daytime yielded an effect of genotype for nest activity, an effect of experimental stage (baseline, shock presentation, or constant conditions), and an interaction for all three behaviors (Table S4). Tukey post hoc comparisons showed statistically significant differences between the different stages of the experiment for all genotypes, indicating that all animals avoided the dark-phase shocks. Visual inspection of the actograms (Fig. 3A and Fig. S4A) as well as analysis of the percent of activity during the light phase (no shocks) and the projected light phase (Fig. 3C and Fig. S4B) revealed that mice with at least one copy of *Bmal1* entrained to the nocturnal fear. In contrast, *Cami-Bmal1*^-/-^ mice avoided the shocks, but this was the result of masking, as they became arhythmic immediately upon release into constant conditions.

**Figure 3.**
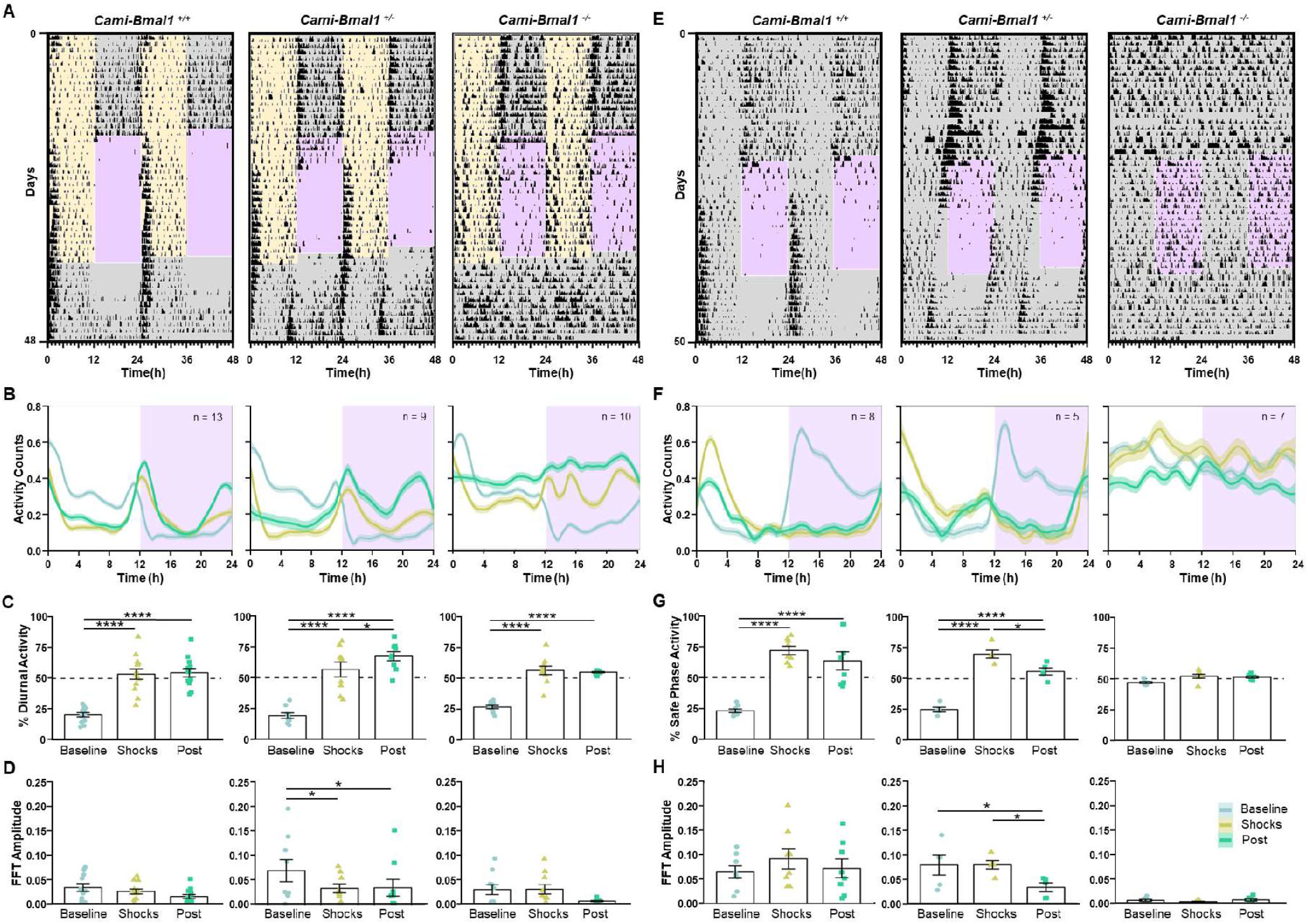
The expression of the clock gene *Bmal1* in the forebrain is necessary for fear entrainment. **(A)** Representative foraging actograms from *Cami-Bmal1*^*+/+*^, *Cami-Bmal1*^*+/-*^ and *Cami-Bmal1*^-/-^ mice subjected to DF. **(B)** Average activity patterns from *Cami-Bmal1*^*+/+*^ (left, n=13), *Cami-Bmal1*^*+/-*^ (center, n=9) and *Cami-Bmal1*^-/-^ mice (right, n=10) in LD subjected to DF. **(C)** Percentage of activity during the daytime or extrapolated daytime across the different experimental stages from the same mice shown in B. **(D)** Fast-Fourier transform (FFT) amplitude across the successive stages from the same mice shown in B. **(E)** Representative foraging actograms from *Cami-Bmal1*^*+/+*^, *Cami-Bmal1*^*+/-*^ and *Cami-Bmal1*^-/-^ mice subjected to a 12-h window of non-cued fear under DD. **(F)** Average activity patterns from *Cami-Bmal1*^*+/+*^ (left, n=8), *Cami-Bmal1*^*+/-*^ (center, n=5) and *Cami-Bmal1*^-/-^ mice (right, n=7) subjected to non-cued fear in DD. **(G)** Percent of activity that took place during the safe phase (window of time without shocks) or extrapolated safe phase across the different experimental stages from the same mice shown in F. **(H)** FFT amplitude across the successive stages from the same mice shown in F. Asterisks indicate statistically significant differences according to Tukey comparisons following LMM analysis: * p<0.05, ** p < 0.01, *** p< 0.001, **** p<0.0001.

To determine how the rhythmicity of each of the three behaviors changed throughout the stages of the experiment, we calculated the amplitude of the fast-Fourier transform (FFT) in a circadian range, which provides an estimate of the robustness of a rhythm. We then fitted a LMM with experimental stage, genotype, and their interaction as fixed factors, and the individual mouse as a random factor. The model yielded an effect of the experimental stage for all three behaviors but no detectable effects of genotype or interaction, revealing the effect of the nocturnal shocks on mice of all genotypes. Most importantly, the apparent lack of any amplitude peak in the circadian frequencies for *Cami-Bmal1*^-/-^ mice confirmed their lack of circadian rhythmicity after their release into constant conditions (Fig. 3D, Fig. S4D, Table S5).

To assess the effect of a dysfunctional molecular clock on fear entrainment under DD conditions, *Cami-Bmal1*^-/-^ mice and their littermate controls were subjected to the non-cued fear protocol in DD. *Cami-Bmal1*^*+/-*^ and *Cami-Bmal1*^*+/+*^ mice were able to avoid the shocks by shifting their activity to the safe time of the day and displayed the expected phase upon release into constant conditions (Fig. 3E, F, Fig. S5A, B, left and center panels). Remarkably, *Cami-Bmal1*^-/-^ mice maintained their arrhythmic pattern during the entire protocol and, during the shock presentation phase, were unable to avoid the shocks by timing their activity to the non-shock phase, suggesting an inability to determine when the aversive stimulus occurred (Fig. 3E, F, Fig. S5A, B, right panels). This conclusion was further supported by the LMM analysis of the percent of activity that took place during the safe time of the day (Fig. 3G, Fig. S5C, Table S6) and of the FFT amplitude (Fig. 3H, Fig. S5D, Table S7). This latter analysis revealed that *Cami-Bmal1*^-/-^ mice lacked amplitude peaks in the circadian frequencies throughout all stages of the experiment.

### An intact molecular circadian clock within the suprachiasmatic nucleus is necessary but not sufficient to sustain fear-entrained circadian rhythms

Our results in *Cami-Bmal1* ^-/-^ mice clearly point to the necessity of an intact molecular circadian clock in the forebrain to sustain cyclic fear entrainment. However, they do not have the regional specificity to determine in which areas of the forebrain clock gene expression is critical for fear entrainment. Specifically, we wondered whether clock gene expression within the SCN central clock is necessary and/or sufficient to sustain fear entrainment of behavioral rhythms. To test this, we injected a Cre-expressing virus into the SCN of *Bmal1*^*F/F*^ mice to delete the *Bmal1* gene specifically from this brain region (Fig. 4A). Mice with off-target injections (no trace of the virus near the SCN) served as controls. As expected, mice lacking *Bmal1* in the SCN (SCN-*Bmal1*^-/-^) became arrhythmic in constant darkness shortly after the viral injection (Figure S6A, B). When subjected to the non-cued fear-entrainment paradigm under DD conditions, SCN-*Bmal1*^-/-^ continued to display an arrhythmic pattern of locomotor activity throughout the protocol, displaying a reduction in the locomotor activity during the shock-presentation portion of the experiment, indicating that these mice were able to sense the aversive nature of the shocks (Fig 4B center panel). In one case, the cyclic fear induced a small degree of activity consolidation in the non-fear phase and a free-running pattern of activity that was coincident with the phase acquired during the shocks (Fig. 4B, right panel). This result suggests that the viral injection spared a small proportion of SCN cells that may have been sufficient for entrainment when forced by the cyclic fearful stimulus presentation. However, we included this mouse in the SCN-*Bmal1*^-/-^ group based on the histological results (see Fig. S6B for an example of histological staining) and the arrhythmicity shown during the baseline stage. The FFT amplitude in the circadian range was quantified at each stage of the protocol as an indicator of the robustness of the rhythm. Control animals showed an overall reduction in amplitude during the post-shock stage relative to the baseline and the shock presentation stage (Fig. 4B, C, Fig. S6C, D left panels, and Table S8). SCN-*Bmal1*^-/-^ showed a lower amplitude FFT throughout all stages of the protocol. Interestingly, for the mouse whose actogram is shown in Fig. 4B right panel, the FFT amplitude increased progressively through the experimental stages, suggesting that a few cells retaining clock genes in the SCN may have been recruited by the fear entrainment. In summary, our results show that a functional circadian clock within the SCN is necessary to sustain fear-entrained circadian rhythms.

**Figure 4.**
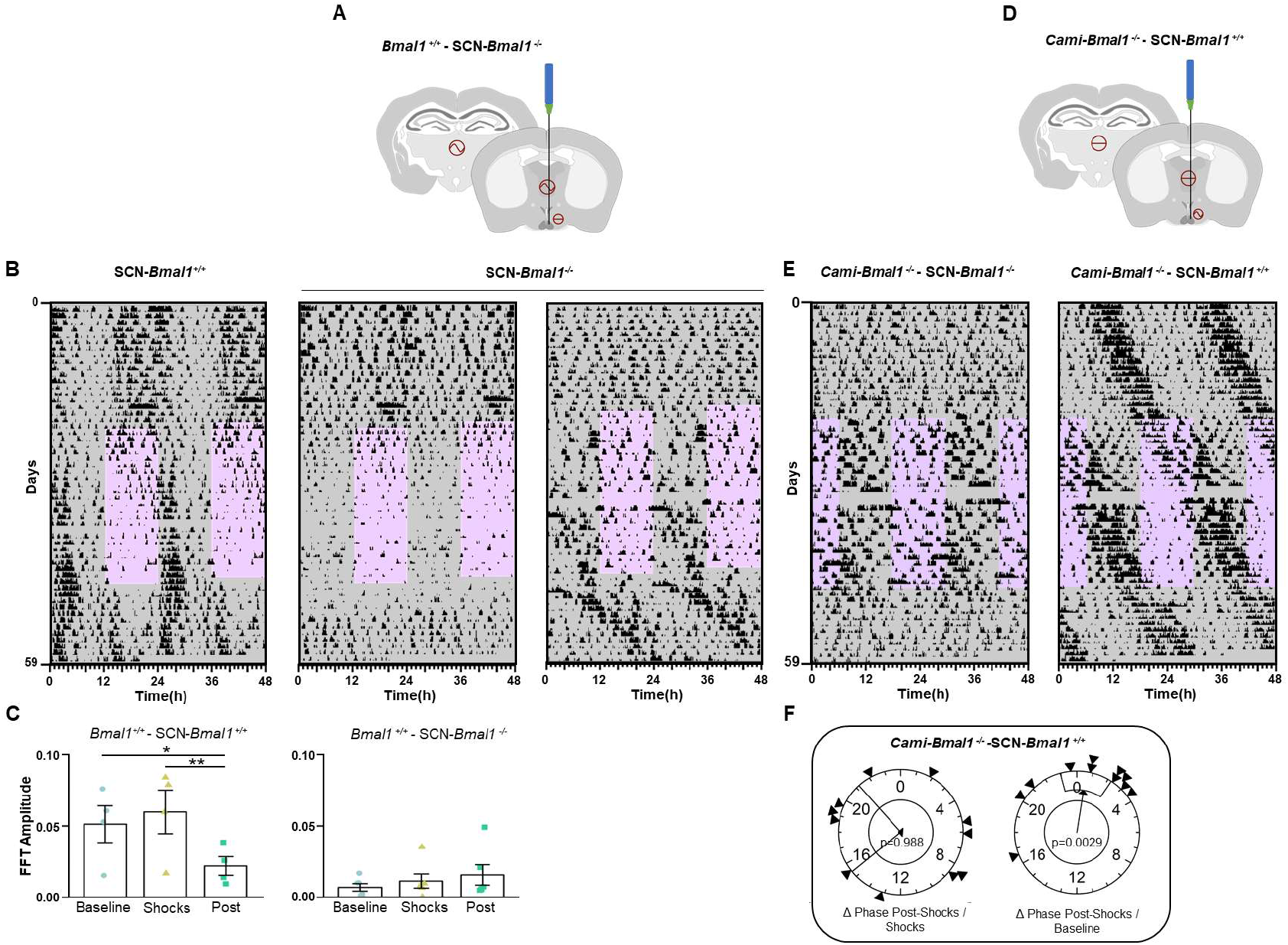
The circadian canonical clock within the SCN is necessary but not sufficient for fear entrainment. **(A)** Schematic representation of the genetic strategy used to knock out *mBmal1* in the SCN. A *Bmal1*^*fx/fx*^ mouse is bilaterally injected with a Cre-expressing AAV in the SCN. **(B)** Representative foraging actograms from an SCN-*Bmal*^*+/+*^ control mouse (left) and two SCN-*Bmal1*^-/-^ mice (center and right) subjected to cyclic fear under DD. **(C)** FFT amplitude across the successive experimental stages from SCN-*Bmal*^*+/+*^ mice (left, n=4) and SCN-*Bmal1*^-/-^ mice (right, n=6). **(D)** Schematic representation of the genetic strategy to rescue *Bmal1* expression in the SCN. A *Cami-Bmal1*^-/-^ mouse is bilaterally injected with Cre-dependent *Bmal1*-expressing AAV. **(E)** Representative foraging actograms from *Cami-Bmal1*^-/-^-SCN-*Bmal1*^-/-^ mouse (left) and *Cami-Bmal1*^-/-^-SCN-*Bmal1*^*+/+*^ mouse (right) subjected to the non-cued fear protocol in DD. **(F)** Rayleigh plots representing the phase of activity onset in the post-shock phase of *Cami-Bmal1*^-/-^-SCN-*Bmal1*^*+/+*^ mice relative to the shock phase (left) or to the baseline phase (right). Asterisks indicate statistically significant differences according to Tukey comparisons following LMM analysis: * p<0.05, ** p < 0.01.

To test whether clock gene expression within the SCN is sufficient to sustain fear-entrained circadian rhythms, we used a complementary approach. We rescued the expression of *Bmal1* in the SCN of mice lacking this gene in the forebrain (*Cami-Bmal1*^-/-^) by injecting a Cre-dependent *Bmal1*-expressing AAV (Fig. 4D).

While control mice (*Cami-Bmal1*^-/-^*-SCN-Bmal1*^-/-^), receiving a control AAV injection or off-target injections, showed arrhythmic patterns of locomotor activity in constant darkness, mice injected with the *Bmal1*-expressing virus within the SCN (*Cami-Bmal1*^-/-^*-SCN-Bmal1*^*+/+*^) recovered the behavioral rhythmicity shortly after the surgery (Fig. S7A, B). When these mice were subjected to the non-cued fear paradigm, they free-ran through the protocol, ignoring the 12-h shock-presentation phase (Fig. 4E). In some individuals, a decrease in foraging and feeding was evident during the shock hours (Fig. 4E, Fig. S7C). Polar plots followed by Rayleigh tests of the rhythm phases showed that the phases of all three behavioral outputs during the post-shock stage were predicted by the phases during the baseline and not by the time of shocks (Fig. 4F, Fig. S7D, Table S9). These results demonstrate a failure of cyclic fear to entrain the rescued rhythms and therefore that a functional circadian clock within the SCN is not sufficient to sustain fear-entrained circadian rhythms.

## Discussion

In the present study, we show that mice living in a more naturalistic environment recreated in the laboratory, where they are required to venture out of their safe nest to obtain food and water, display nocturnal food-seeking and feeding behavior. However, the chronic application of a nocturnal aversive stimulus in the foraging area leads to a shift in foraging and feeding behavior to daytime. This shift results from cyclic fear entrainment of a circadian oscillator, similar to what occurs in rats (*7*). Our results suggest that cyclic fear is a potent non-photic entraining environmental cycle or *zeitgeber* for the circadian system, capable of dramatically shifting 24-h patterns of overt behavior. Furthermore, we demonstrate that this fear-entrained oscillator relies on the canonical circadian molecular clock. Using conditional KO strategies, we also show that an intact molecular clock within the central circadian clock located in the SCN is necessary but not sufficient to sustain fear entrainment.

Virtually all organisms rely on circadian systems to predict 24-h cyclic events in nature. Internal biological clocks provide a mechanism by which organisms can anticipate these events with changes in physiology and behavior. Because circadian clocks have periods that differ from 24 h, they require entrainment by 24-h environmental cycles. Throughout evolution, the LD cycle has been selected as a highly reliable cycle conveying solar time, and the circadian clocks of most organisms are entrained by it. However, animals live in complex temporal environments, and their circadian system can be entrained by non-photic zeitgebers as well (*13, 14*). A classic example of this non-photic entrainment is time-restricted feeding, which leads to activity in anticipation of feeding events and is the result of entrainment by a food-entrainable oscillator(s) (FEO) (*15, 16*). Here, we show that cyclic fear is similarly effective in entraining circadian rhythms. Given that the ability of animals to perceive and respond appropriately to threats directly affects individual survival, it is logical that cyclic fear acts as a strong zeitgeber. This adaptive response should reliably reduce the negative fitness consequences of encountering recurring dangers, such as predators, in the wild.

Despite the discovery of the FEO over half a century ago, the basic molecular mechanisms behind this clock and its precise location in the brain or body have remained remarkably elusive (*10, 17*). Animals devoid of a canonical circadian molecular clock, as well as those with complete lesions of the SCN, can still entrain to restricted food access. In contrast, rats with lesions of the SCN are unable to entrain to nocturnal fear (*7*). In this study, we further demonstrate that a functional molecular clock within the SCN is necessary for fear entrainment. This finding potentially suggests that the SCN can be entrained by cyclic fear in a similar manner to how it is entrained by the LD cycle. However, this is likely not the case for two reasons. First, we show that when mice entrain to nocturnal fear, clock gene expression in the SCN remains synchronized to the LD cycle and not to the timing of fear. Second, animals with a functional SCN, but non-functional clocks in the rest of the forebrain fail to entrain to cyclic fear. This latter result suggests that other brain centers that rely on a canonical molecular circadian clock can entrain to cyclic fear. However, neither the amygdala nor the dentate gyrus exhibits synchronized clock gene expression with cyclic fear.

The necessity, but not sufficiency, of the SCN for animals to entrain to cyclic fear suggests that its role is to convey phase information about the LD cycle. However, we also show that, remarkably, animals cannot entrain to fear without a functional SCN, even under constant darkness (DD) conditions. Thus, the SCN may provide an internal circadian phase reference, enabling the scheduling of foraging, feeding, and nest activity to avoid a circadian time in which a threat is present.

The discovery of circadian oscillators outside the SCN about two decades ago challenged the classic layout of a central circadian clock governing overt circadian rhythms. It soon became clear that clocks downstream of the SCN played an important role as subordinate clocks, timing local rhythmic outputs such as enzymatic pathways in the liver or glucocorticoid production in the adrenal gland. This layout was further complicated by examples where these subordinate clocks could independently synchronize to external cycles and override central control by the SCN. For instance, restricted food access during the light phase in mice entrains the liver clock and extra-SCN oscillators in the brain but does not entrain the SCN (*18-20*). This change in the configuration of the ensemble of circadian oscillators is associated with daytime activity anticipating food arrival, indicating that the typical hierarchy in which the SCN is the leading oscillator timing daily activity can be altered, allowing other oscillators to take control. Similarly, our results show that cyclic aversive stimuli can entrain overt patterns of behavior, clearly indicating that fear-coding centers are an integral component of the circadian system, alongside the liver, the retina, and the central SCN pacemaker.

This notion may have significant implications for understanding symptoms of fear and anxiety disorders, which are often associated with sleep and circadian disruptions, particularly in patients suffering from post-traumatic stress disorder (PTSD). Our results show that cyclic aversive stimuli can time-stamp the circadian system, leading to changes in the timing of behavior that persist even after the aversive stimulus is removed, leaving only its fear engram remains. This supports the interpretation that sleep and circadian disorders associated with PTSD could be the result of a circadian phase that cyclically reenacts a fear sensation.

## Author contributions

Conceptualization: ILB, MBH, HOD

Methodology: ILB, MBH, LESL, LC, AN, HOD

Investigation: ILB, MBH, LESL, VYZ, LC, AN, JL, HOD

Funding acquisition: ILB, JST, HOD

Project administration: HOD

Writing – original draft: ILB, HOD

Writing – review & editing: ILB, MBH, LESL, VYZ, LC, AN, JL, JST, HOD

## Data and materials availability

All data, code, and materials used in the analysis will be available upon request from the authors.

## Supplementary Materials

Materials and Methods

Figs. S1 to S7

Tables S1 to S9

