## Supplementary material for "The circadian molecular clock in the suprachiasmatic nucleus is necessary but not sufficient for fear entrainment in the mouse"

**The PDF file includes:**

Materials and Methods
Figs. S1 to S7
Tables S1 to S9

### Materials and Methods

Animals

C57BL/6J male and female mice purchased from The Jackson Laboratory were used for all experiments carried out with wild type mice. The light-fear (LF), dark-fear (DF), cued and non-cued fear in constant darkness experiments with wild type mice were done with male mice only. These experiments were replicated with both females and males, and all remaining experiments had approximately 50% females. For none of the reported behavioral or histological assays did sex have an effect on the outcome variables, and results represent the aggregated data of males and females. *Cami-Bmal1* mice were generated as previously described (*1*). Briefly, *Camk2a::iCre*BAC mice (*CamiCre*+/-) (MGI:2181426), were crossed to *Bmal1*$^{fx/fx}$ mice (The Jackson Laboratory Stock number 007668) to produce *CamiCre*$^{+/-}$*;Bmal1*$^{fx/+}$. These animals were backcrossed to *Bmal1*$^{fx/fx}$ to produce *CamiCre*$^{+/-}$*;Bmal1*$^{fx/fx}$ *(Cami-Bmal1*$^{-/-}$), *CamiCre*$^{+/-}$*;Bmal1*$^{fx/+}$ *(Cami-Bmal1*$^{+/-}$) and *CamiCre*$^{-/-}$*;Bmal1*$^{fx/fx}$ *(Cami-Bmal1*$^{+/+}$).

To induce the deletion of the *Bmal1* gene specifically in the adult SCN, *Bmal1*$^{fx/fx}$ adult mice were injected bilaterally at the SCN with a Cre and GFP-expressing AAV (AAV2/1-Ef1a-Gfp-Cre).

To rescue the expression of *Bmal1* in the SCN of mice lacking its expression in the forebrain, *CamiCre-;Bmal1*$^{fx/fx}$ mice were injected with a Cre-dependent *Bmal1*-expressing AAV (AAV2/1-Ef1a-DIO-Bmal1). The viruses were produced in the Vision Core Lab at the University of Washington. The *Bmal1*-expressing plasmid was custom made by VectorBuilder (Chicago, IL, USA). Targeted viral injections to the SCN were performed aseptically while mice were head-fixed on a stereotaxic device and anesthetized with isoflurane. The injection coordinates were: anteroposterior -0.5 mm, mediolateral ± 0.25 mm, and dorsoventral -5.65 mm. Viruses were loaded into pulled glass capillary needles that were backfilled with biologically inert Perfluro-compound (FC-770) and administered using a Nanoject II (Drummond Scientific) 500 nL of virus was injected at a working concentration of $10^{12}$ particles/ml to each hemisphere of the SCN.

Cyclic fear paradigm

Animals were housed in regular (19 x 40 x 18 cm WxLxD) mouse cages under a 12:12 LD cycle unless otherwise indicated. For fear-entrainment experiments each mouse was singly housed in a fear conditioning chamber. The chamber is composed of a nesting area (11x21x20 cm WxLxH) containing corncob bedding connected to a “foraging area” (10x21x20 cm) that provided ad libitum access to food and water (Fig. S1A). The floor of the foraging area consisted of a foot-shock grid that was connected to an Arduino-controlled (Arduino Uno; New York, NY) shocker that could be programed to deliver shocks with any temporal structure. Two independent IR detectors recorded the activity within the nesting and foraging areas, respectively, and a laser bin detected nose-pokes into the food container. Thus, we continuously recorded nest activity, foraging (activity within the foraging area), and feeding for each individual animal.

The basic fear-entrainment protocol consisted of three different phases. First, a baseline phase (~14 days) in which the mice were allowed to familiarize with the new environment without receiving any aversive stimulus. Second, the shocks phase (10-15 days), in which the aversive stimulus, three 2-mAmp foot shocks per hour randomly distributed, was incorporated in a daily 12-h window in a phase-specific manner (see below). During the final post-shock phase (7-10 days), mice are released into constant darkness conditions and the aversive stimulus was removed. This free-running phase was necessary to evaluate whether rhythmic activity as the result of the time-specific shocks was the result of circadian entrainment.

Light and dark fear entrainment protocol

In mice subjected to footshocks during the light phase (light fear = LF) protocol, the 12-h window of shock presentation was paired to the 12-h light phase of the LD cycle. For mice subjected to footshocks during the dark phase (dark fear = DF), the stimulus was presented in the opposite phase, paired with the 12-h dark phase.

Cued and non-cued constant darkness fear entrainment protocol

Mice were first placed in the fear chamber under an LD cycle for a minimum of 7 days and were subsequently transferred into DD (dim red light of <2-lux intensity). After a 14-day-long baseline phase, one group of animals received a 12-h window of shocks with the same temporal structure as above (non-cued fear group) whereas the second group received the same temporal pattern of shocks but each shock was preceded by a 20-sec 4 kHz 75dB tone that served as conditioned stimulus (cued fear group).

In situ hybridization

Mice were subjected to a baseline phase (10 days) followed by either LF or DF phases (15 days) (as described in LD protocol above). During the last day of shock exposures, mice were euthanized, and their brains collected and frozen every 4 hours for 24 hours (ZT 2, 6, 10, 14, 18, 22). Frozen brains were sliced in 16um-thick coronal slices by using a cryostat and mounted on vectabonded slides. *In situ* hybridization for *Per1* and *Bmal1* genes was conducted as previously described (de la Iglesia, 2007; Han et al., 2012). Autoradiographic images were generated by exposing slides to Ultramax film (Kodak, Rochester, NY). Images were scanned at high resolution, and hybridization intensities were determined with ImageJ software (National Institutes of Health, Bethesda, MD).

Immunohistochemistry

Brain tissue for immunohistochemistry (IHC) was harvested at ZT 9-11 under LD 12:12 conditions. Briefly, mice were anesthetized with Isoflurane and perfused transcardially using 0.1M phosphate buffer followed by 4% paraformaldehyde in 0.1M phosphate buffer. Brains were removed and postfixed overnight at 4°C in 4% paraformaldehyde in 0.1M phosphate buffer and then incubated at 4°C in 30% sucrose/phosphate buffer overnight.

Cryostat 30-um coronal sections were collected into 3 alternate sets representing the whole rostro-caudal extent of the SCN and used for free-floating IHC. Briefly, the slices were incubated in 0.4% tween 20 in 0.01M phosphate buffered saline (PBST) for 15 minutes to permeabilize tissue and then blocked in 5% donkey serum in PBST for an hour. After a quick rinse in 0.04% PBST, the slices were incubated in BMAL1 (1:1000, Novus Biological cat# NB100-2288) primary antibody solution at 4ºC overnight. Then, the slices were washed four times, 10 min each in 0.04% PBST and incubated in Alexa 594 Donkey anti-Rabbit secondary antibody diluted in 0.04% PBST (1:300, Invitrogen cat# A21207) for 2 hours at room temperature. After the secondary antibody incubation, the slices were washed four times, 5 min each with 0.04% PBST, and mounted into glass slides and cover-slipped with DAPI Fluoromount-G (Southern Biotech, Birmingham, AL).

Images were taken using a Leica TCS SP5 II laser scanning confocal microscope (20X objective) in 1-um Z-stacks using identical capture settings for every slice.

Statistical analysis of behavioral outputs

Average daily activity patterns (or "waveforms") of the different outputs measured were generated using 7 days of recording for each stage in the fear entrainment protocol (the last 7 days for baseline and shocks phases, and the first 7 days of post-shocks phase). The percentage of activity performed during the light phase for experiments carried under LD conditions, and the

percentage of activity displayed during the safe phase (no-shocks) in DD conditions were calculated using El Temps software (University of Barcelona, Spain) . The onset of activity was calculated using El Temps Software and those phases were evaluated using Raleigh test.
To evaluate the robustness of the rhythms in the behavioral outputs measured, the relative power from the FFT for the peak within the circadian range (20-28hs) was measured for each mouse (Clocklab Analysis, Actimetrics) and used for statistical analysis.
Linear mixed-effects models (LMM) were used to analyze differences in the percent of locomotor and feeding activity at specific phases, and in FFT power across the protocol stages using the lme4 package for R (*2*). Statistical significance for LMM factors was calculated through a Type III analysis using Satterthwaite's method using the lmerTest package (*3*).Post-hoc Tukey comparisons within groups were performed using the emmeans package (*4*).
Expression of genes measured by in-situ hybridization were fit to a cosinor model, and differences in fit parameters were analyzed through Wald tests using the cosinor package for R (*5*).
Group-wise average activity patterns were calculated and plotted using a locally estimated scatterplot smoothing (LOESS) using tools from the tidyverse package for R (*6*).

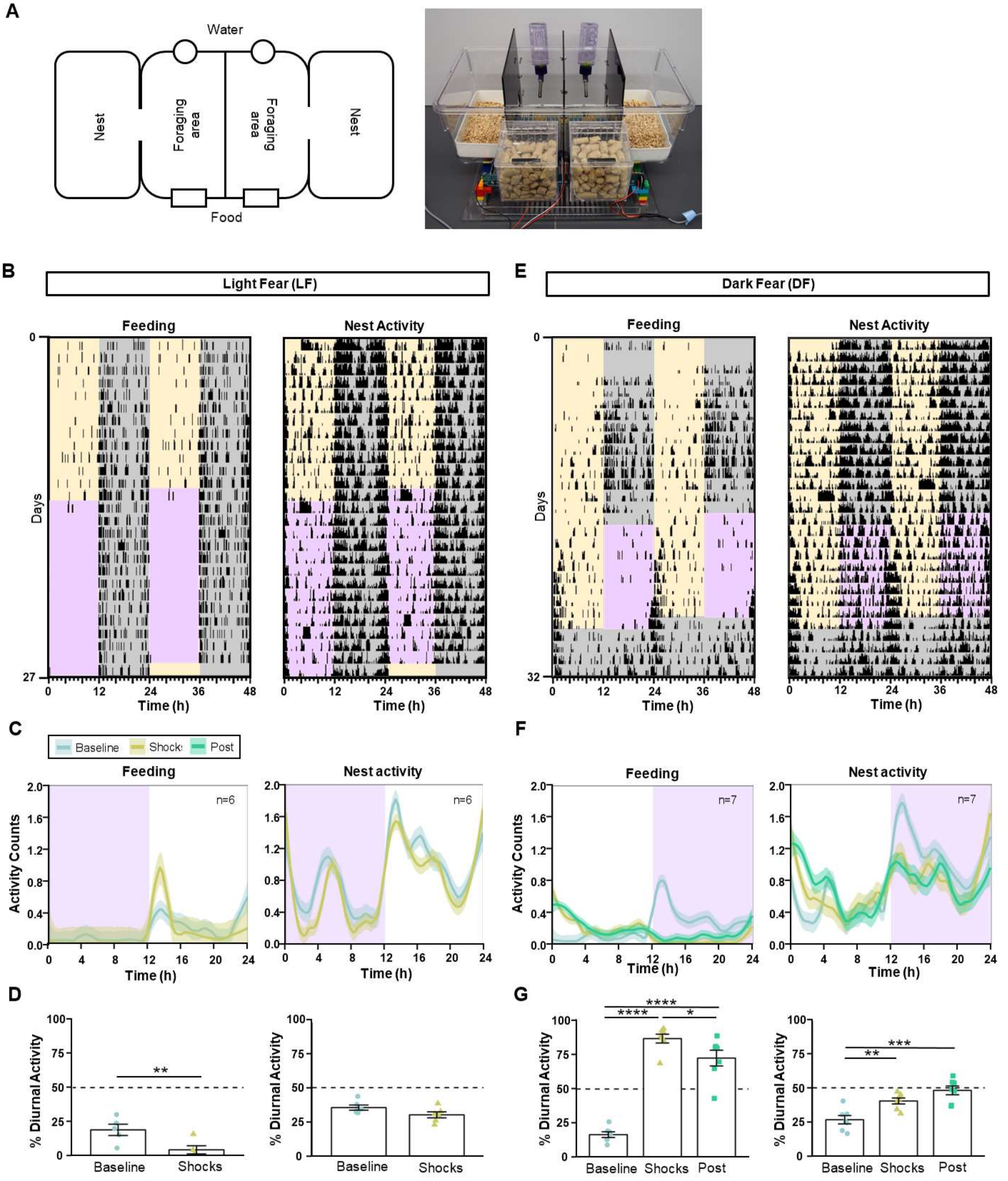


**Figure S1. Cyclic fear entrains a circadian oscillator under a light-dark cycle. (A)** Custom-built cages to study fear entrainment. Top: Schematic of a cage where two mice are individually housed. Each mouse has access to a safe nest cage and needs to get all the food and water from a foraging area. The floor of the foraging area can deliver footshocks with any temporal pattern.

The shocking grid is shared by the two mice but each mouse's setup is separated by an opaque divider. Bottom: Photograph of the fear-entrainment setup. A nose poke detector continuously records feeding from each feeder. Individual IR sensors on the cage lid (not displayed) monitor the activity of each mouse independently in the nest and foraging areas. **(B)** Representative feeding and nest activity actograms from mice in LD subjected to LF (left) or DF (right). Yellow and grey shading respectively represents the light and dark phases of a 12:12 LD cycle. Purple shading represents the 12-h window of time at which 3 footshocks/hour randomly distributed over time were presented. **(C)** Average feeding and nest activity patterns from mice under an LD cycle subjected LF (left, n=6) or DF (right, n=7). **(D)** Percent of activity that took place during the daytime or extrapolated daytime across the different experimental stages from the same mice shown in B. Bars represent mean ± SEM. *P < 0.05, **P < 0.01, ***P < 0.001. Symbols as in Fig. 1.

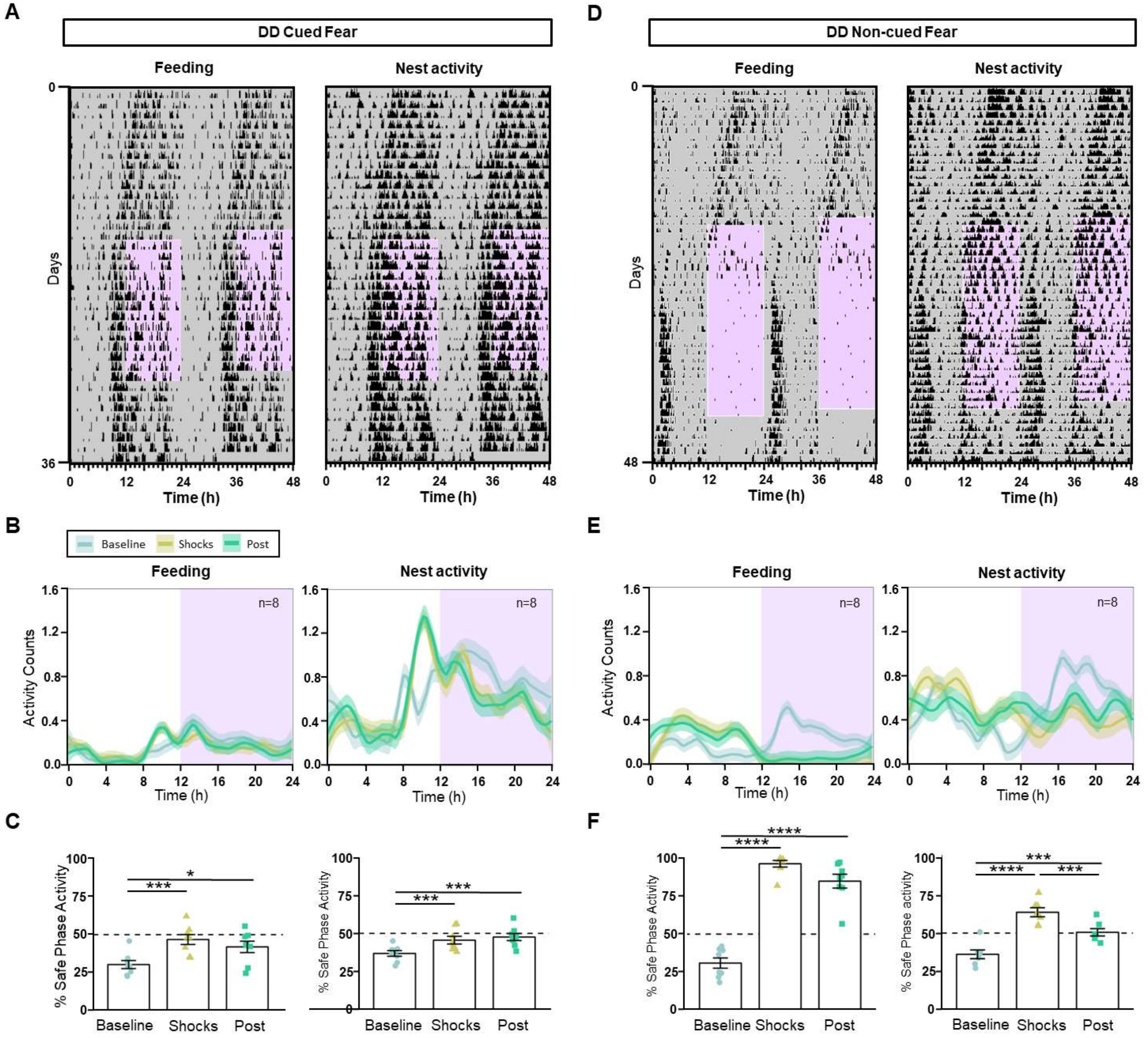


**Figure S2. Cyclic fear entrains a circadian oscillator under constant darkness. (A, D)** Representative feeding and nest activity actograms from mice in DD subjected to either cued (A) or non-cued (D) cyclic fear. Purple shading represents the 12-h window of time at which 3 footshocks/hour randomly distributed over time were presented either preceded by an auditory signal (A, cued) or not (D, non-cued). **(B, E)** Average feeding and nest activity patterns from mice in DD subjected to cued (B, n=8) or non-cued fear (F, n=8). **(C, F)** Percent of activity that took place during the safe phase (window of time without shocks) or extrapolated safe phase across the different experimental stages from the same mice shown in B, F. *P < 0.05, ***P < 0.001. Symbols as in Fig. 1.

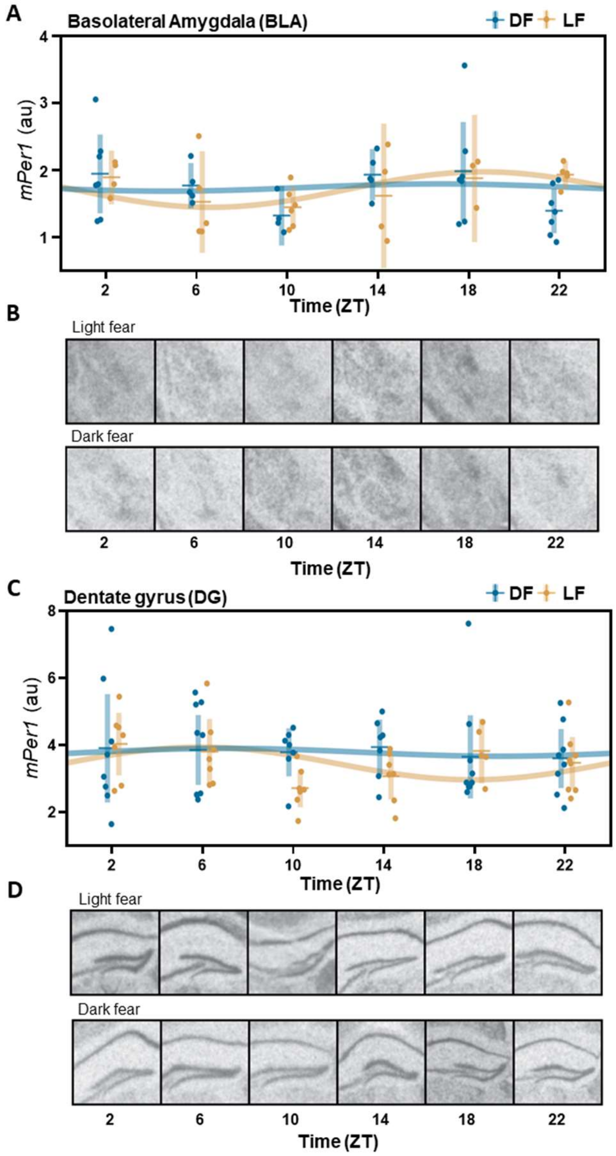


**Figure S3**. m***Per1*** **expression in in the basolateral amygdala and dentate gyrus**. (A, C ) Daily pattern of *mPer1* mRNA expression in mice housed under a 12:12 LD cycle and subjected to or DF. Each dot represents an individual mouse, horizontal and vertical lines respectively represent the mean and SEM. Cosinor analysis failed to detect any oscillation in any of the regions or groups. (B, D) Representative autoradiographs of coronal brain sections at the level of the basolateral amygdala (B) and the dentate gyrus of the hippocampus, hybridized with a radioactive probe for mPer1mRNA detection.

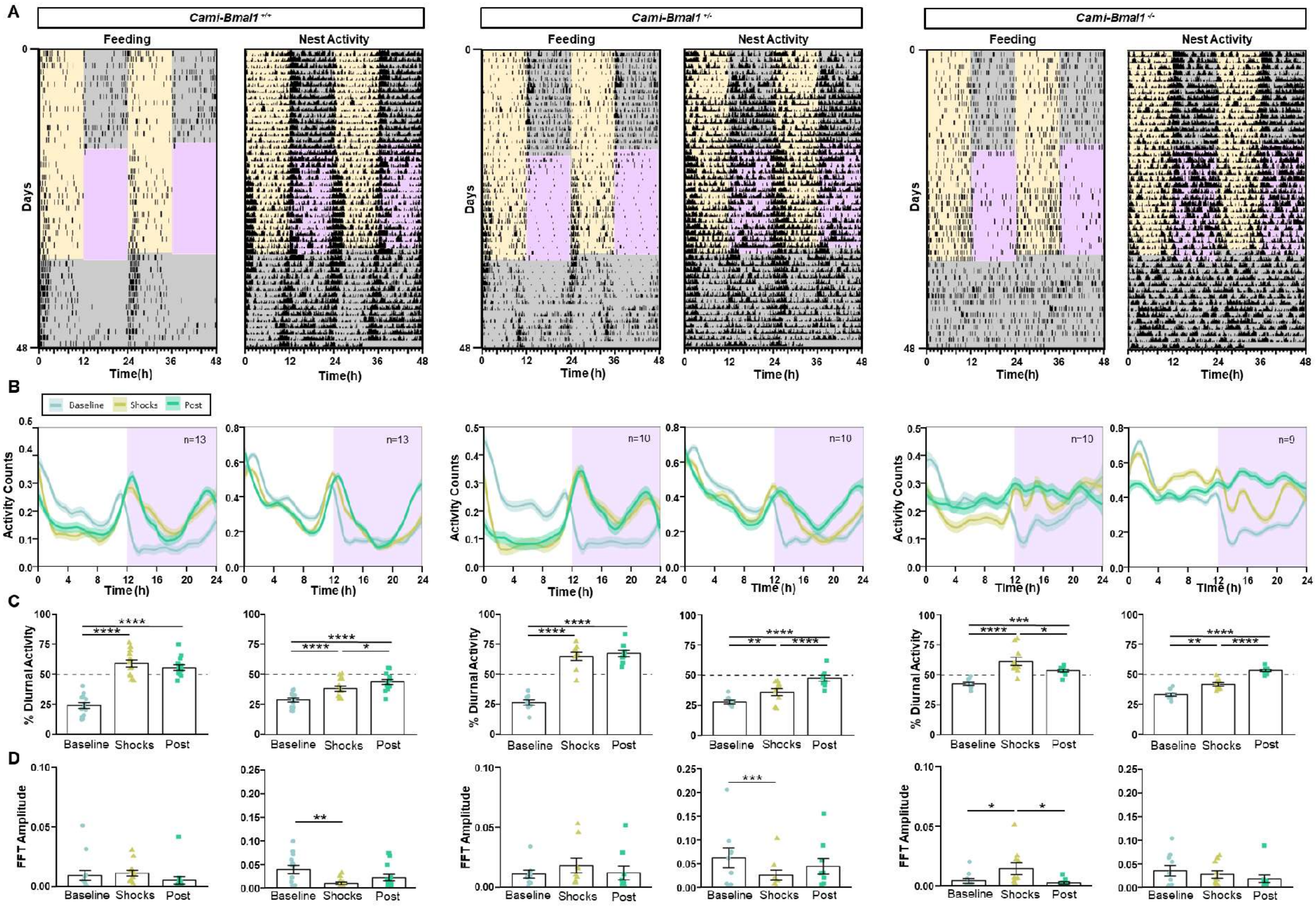
A
Cami-Bmal1 +/+
Cami-Bmal1 +/-
Cami-Bmal1 -/-
Feeding
Nest Activity
Days
Time(h)
B
Baseline
Shocks
Post
Activity Counts
n=13
n=10
n=9
Time (h)
C
% Diurnal Activity
Baseline
Shocks
Post
D
FFT Amplitude

**Figure S4. The expression of the clock gene *Bmal1* in the forebrain is necessary for fear entrainment.** (A) Representative feeding and nest activity actograms from Cami-Bmal1$^{+/+}$, Cami-Bmal1$^{+/-}$ and Cami-Bmal1$^{-/-}$ mice subjected to DF. (B) Average activity patterns from Cami-Bmal1$^{+/+}$ (left, n=13), Cami-Bmal1$^{+/-}$ (center, n=9) and Cami-Bmal1$^{-/-}$ mice (right, n=10) subjected to DF. (C) Percentage of activity during the daytime or extrapolated daytime across the different experimental stages from the same mice shown in B. (D) Fast-Fourier transform (FFT) Amplitude across the successive stages from the same mice shown in B. *P < 0.05, **P < 0.01, ***P < 0.001. Symbols as in Fig. 1.

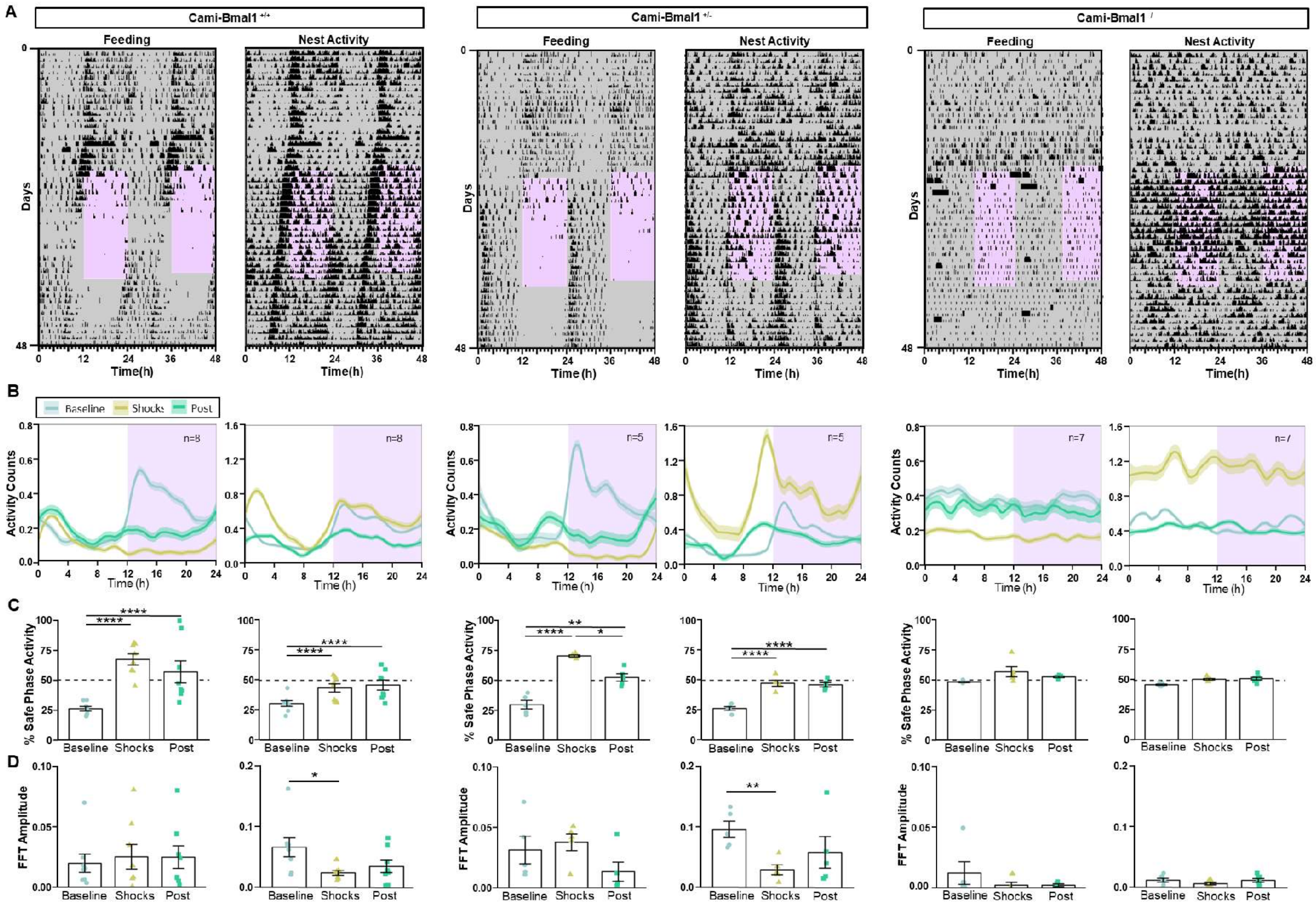

A
Cami-Bmal1 +/+
Cami-Bmal1 +/-
Feeding
Nest Activity
Days
Time(h)
B
Baseline
Shocks
Post
Activity Counts
Time (h)
n=8
n=5
n=7
C
% Safe Phase Activity
Baseline
Shocks
Post
D
FFT Amplitude

**Figure S5. The expression of the clock gene *Bmal1* in the forebrain is necessary for fear entrainment. (A)** Representative feeding and nest activity actograms from Cami-Bmal1$^{+/+}$, Cami-Bmal1$^{+/-}$ and Cami-Bmal1$^{-/-}$ mice subjected to a non-cued fear protocol in DD. **(B)** Average activity patterns from Cami-Bmal1$^{+/+}$ (left, n=8), Cami-Bmal1$^{+/-}$ (center, n=5) and Cami-Bmal1$^{-/-}$ mice (right, n=7) subjected to a non-cued fear protocol in DD. **(C)** Percent of activity that took place during the safe phase (window of time without shocks) or extrapolated safe phase across the different experimental stages from the same mice shown in F. **(D)** FFT amplitude across the successive stages from the same mice shown in C. *P < 0.05, **P < 0.01, ***P < 0.001. Symbols as in Fig. 1.

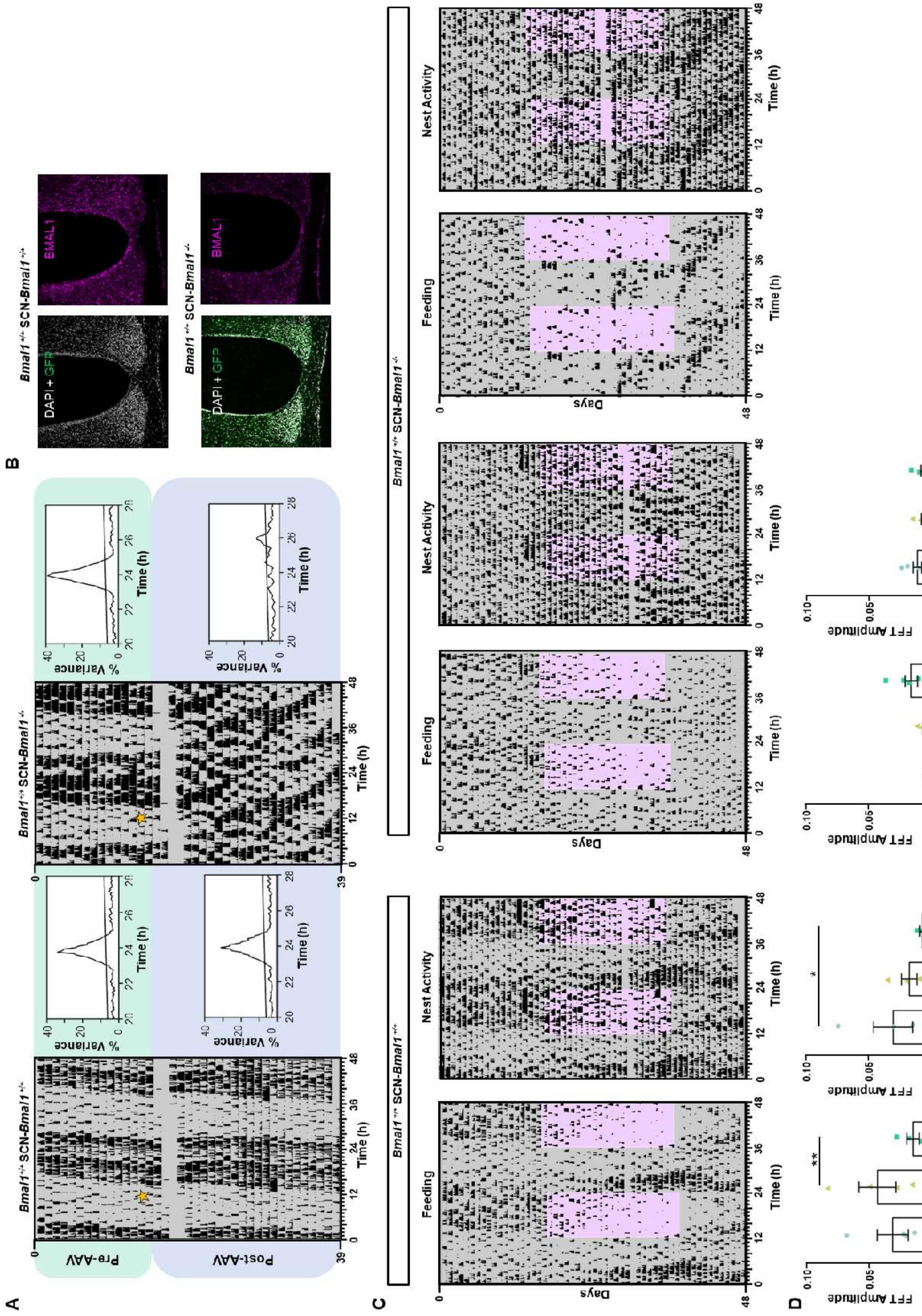
A
Bmal1+/+ SCN-Bmal1+/+
Bmal1+/+ SCN-Bmal1-/-
Pre-AAV
Post-AAV
% Variance
Time (h)
B
Bmal1+/+ SCN-Bmal1+/+
Bmal1+/+ SCN-Bmal1-/-
DAPI + GFP
BMAL1
C
Bmal1+/+ SCN-Bmal1+/+
Bmal1+/+ SCN-Bmal1-/-
Feeding
Nest Activity
Days
Time (h)
D
FFT Amplitude
Baseline
Shocks
Post

**Figure S6. The circadian canonical clock within the SCN is necessary for fear entrainment. (A)** Actograms and periodograms of cage locomotor activity of two representative mice before and after they received an injection of a Cre-expressing AAV outside (left, SCN-*Bmal1*$^{+/+}$) or within (right, SCN-*Bma1l*$^{-/-}$) the SCN. **(B)** Coronal sections at the level of the SCN from two representative mice displaying fluorescence for GFP (expressed by the Cre-expressing AAV) and immunostained for BMAL1. **(C)** Representative feeding and nest activity actograms from a SCN-*Bmal1*$^{+/+}$ control mouse (left) and two SCN-*Bmal1*$^{-/-}$ mice (center and right) subjected to non-cued fear in DD (same individuals shown in fig. 4B). **(D)** FFT Amplitude across the successive experimental stages from SCN- *Bmal*$^{+/+}$ mice (left, n=4) and SCN-*Bmal1*$^{-/-}$ mice (right, n=6). *P < 0.05, **P < 0.01. Symbols as in Fig. 1.

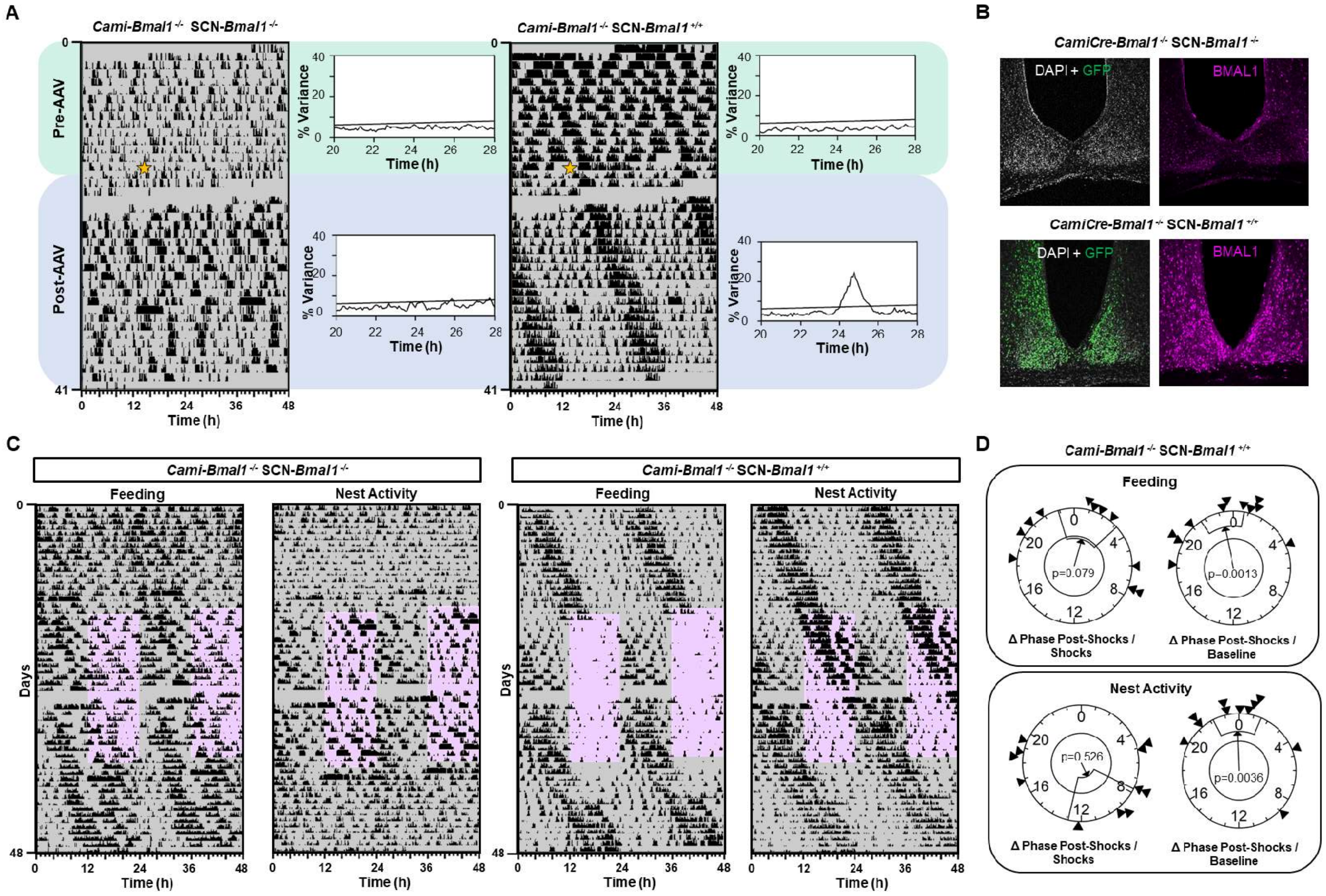
A
Cami-Bmal1 -/- SCN-Bmal1 -/-
Cami-Bmal1 -/- SCN-Bmal1 +/+
Pre-AAV
Post-AAV
% Variance
Time (h)
B
CamiCre-Bmal1 -/- SCN-Bmal1 -/-
CamiCre-Bmal1 -/- SCN-Bmal1 +/+
DAPI + GFP
BMAL1
C
Cami-Bmal1 -/- SCN-Bmal1 -/-
Cami-Bmal1 -/- SCN-Bmal1 +/+
Feeding
Nest Activity
Days
Time (h)
D
Cami-Bmal1 -/- SCN-Bmal1 +/+
Feeding
p=0.079
p=0.0013
Nest Activity
p=0.526
p=0.0036
Δ Phase Post-Shocks / Shocks
Δ Phase Post-Shocks / Baseline

**Figure S7. The circadian canonical clock within the SCN is not sufficient for fear entrainment. (A)** Actograms and periodograms of cage locomotor activity of two representative mice before and after they received an injection of a *Bmal1*-expressing AAV outside (left, Cami-Bmal1$^{-/-}$ SCN-Bmal1$^{-/-}$) or within (right, , Cami-Bmal1$^{-/-}$ SCN-Bmal1$^{+/+}$) the SCN. **(B)** Coronal sections at the level of the SCN of two representative mice displaying fluorescence for GFP (expressed by the *Bmal1*-expressing AAV) and immunostained for BMAL1. **(C)** Representative feeding and nest activity actograms from *Cami-Bmal1*$^{-/-}$-SCN-*Bmal1*$^{-/-}$ mouse (left) and *Cami-Bmal1*$^{-/-}$-SCN-*Bmal1*$^{+/+}$ mouse (right) subjected to non-cued fear protocol in DD (same individuals shown in fig. 4E). **(D)** Rayleigh plots representing the phase of the feeding and nest activity onset in the post-shock phase of *Cami-Bmal1*$^{-/-}$-SCN-*Bmal1*$^{+/+}$ mice relative to the shock phase (left) or to the baseline phase (right). Symbols as in Fig. 1.

**Table S1.** Results for the linear models with mixed effects of the percent of diurnal or safe time activity in mice entrained to fear under LD or DD conditions. Statistics correspond to Figs. 1C, G and Figs S1D, G and Fig. S2C, F.

| Experiment | Behavior | n | Stage | Mean % of diurnal or safe-time activity | SEM | 95% CI | | | Tukey | LMM-AOV |
|---|---|---|---|---|---|---|---|---|---|---|
| LF | Feeding | 6 | Baseline | 19.3 | 4.1 | 7.9 | - | 30.7 | a | **0.0079** |
| | | | Shocks | 4.8 | 3 | -3.4 | - | 13 | b | |
| | Foraging | 6 | Baseline | 21.4 | 3.3 | 12.9 | - | 29.9 | a | **0.0006** |
| | | | Shocks | 5.4 | 1.5 | 1.4 | - | 9.3 | b | |
| | Nest activity | 6 | Baseline | 35.6 | 1.9 | 30.8 | - | 40.3 | a | 0.0999 |
| | | | Shocks | 30.3 | 2.2 | 24.7 | - | 35.9 | a | |
| DF | Feeding | 7 | Baseline | 17.1 | 2.1 | 12 | - | 22.1 | a | **9.78E-09** |
| | | | Shocks | 86.4 | 3.2 | 78.6 | - | 94.1 | b | |
| | | | Post | 72.3 | 5.7 | 58.5 | - | 86.2 | c | |
| | Foraging | 7 | Baseline | 23.3 | 3.7 | 14.3 | - | 32.2 | a | **1.23E-08** |
| | | | Shocks | 80.5 | 3.3 | 72.4 | - | 88.6 | b | |
| | | | Post | 68.1 | 4.7 | 56.7 | - | 79.5 | b | |
| | Nest activity | 7 | Baseline | 27.1 | 3 | 19.9 | - | 34.4 | a | **0.0003** |
| | | | Shocks | 40.6 | 2.1 | 35.3 | - | 45.8 | b | |
| | | | Post | 48.2 | 3.1 | 40.5 | - | 55.8 | b | |
| Cued Fear | Feeding | 8 | Baseline | 30.2 | 2.7 | 23.8 | - | 36.5 | a | **0.0007** |
| | | | Shocks | 46.8 | 3.3 | 39 | - | 54.5 | b | |
| | | | Post | 41.9 | 3.8 | 32.9 | - | 50.8 | b | |
| | Foraging | 8 | Baseline | 33.2 | 2.8 | 26.7 | - | 39.8 | a | **0.0011** |
| | | | Shocks | 43.4 | 3 | 36.2 | - | 50.6 | b | |
| | | | Post | 41.2 | 2.8 | 34.6 | - | 47.9 | b | |
| | Nest activity | 8 | Baseline | 36.9 | 1.8 | 32.5 | - | 41.3 | a | **0.0001** |
| | | | Shocks | 45.6 | 2.5 | 39.6 | - | 51.5 | b | |
| | | | Post | 47.6 | 2.3 | 42.2 | - | 53 | b | |
| Non-cued Fear | Feeding | 8 | Baseline | 30.9 | 3.4 | 22.9 | - | 38.8 | a | **2.01E-11** |
| | | | Shocks | 96.4 | 2.2 | 91.3 | - | 101.6 | b | |
| | | | Post | 84.9 | 4.6 | 74.1 | - | 95.7 | b | |
| | Foraging | 8 | Baseline | 30.5 | 2.2 | 25.2 | - | 35.8 | a | **3.75E-11** |
| | | | Shocks | 94.6 | 3.1 | 87.3 | - | 101.9 | b | |
| | | | Post | 78.7 | 3.7 | 69.9 | - | 87.5 | c | |
| | Nest activity | 8 | Baseline | 36.2 | 2.8 | 29.3 | - | 43.2 | a | **5.33E-07** |
| | | | Shocks | 63.5 | 2.9 | 56.3 | - | 70.7 | b | |
| | | | Post | 50.5 | 2.4 | 44.7 | - | 56.3 | c | |

**Table S2.** Results for Rayleigh statistics on the phase of mice entrained to fear under LD or DD conditions. Statistics correspond to Figs. 1D, H.

| | | | | Rayleigh statistics | | | | |
|---|---|---|---|---|---|---|---|---|
| **Condition** | **Treatment** | **Stage** | **n** | **r** | **p** | **Phase (h)** | **Fiduciary Limits** | **SEM** |
| LD | LF | Baseline | 6 | 0.99 | 0.0003 | 12.0 | 11.93 - 12.15 | 0.05 |
| | | Shocks | 6 | 0.99 | 0.0003 | 12.1 | 12.04 - 12.18 | 0.03 |
| | | - | - | - | - | - | - | - |
| | DF | Baseline | 7 | 0.99 | 0.0001 | 11.9 | 11.66 - 12.18 | 0.13 |
| | | Shocks | 7 | 0.92 | 0.0006 | 0.4 | 23.36 - 1.53 | 0.55 |
| | | Post | 7 | 0.99 | 0.0001 | 23.0 | 22.55 - 23.42 | 0.22 |
| DD | Cued Fear | Baseline | 8 | 0.87 | 0.0007 | 10.1 | 8.83 - 11.40 | 0.68 |
| | | Shocks | 8 | 0.87 | 0.0008 | 9.5 | 8.16 - 10.81 | 0.70 |
| | | Post | 8 | 0.86 | 0.0009 | 9.4 | 7.99 - 10.71 | 0.71 |
| | Non-cued Fear | Baseline | 8 | 0.97 | <0.0001 | 14.2 | 13.54 - 14.81 | 0.33 |
| | | Shocks | 8 | 0.85 | 0.001 | 2.1 | 0.77 - 3.52 | 0.72 |
| | | Post | 8 | 0.82 | 0.002 | 2.0 | 0.47 - 3.54 | 0.81 |

**Table S3.** Cosinor analysis for w4-h clock gene expression in the SCN, dentate gyrus (DG) and basolateral amygdala (BLA)

| | **DF** | | | | **LF** | | | | **Wald tests** | |
|---|---|---|---|---|---|---|---|---|---|---|
| ***SCN mPer1*** | **Estimate** | **CI 95%** | | | **Estimate** | **CI 95%** | | | **Mean Difference** | **p-value** |
| Mean (au) | 3.42 | 3.11 | - | 3.73 | 3.59 | 3.31 | - | 4.21 | - | - |
| Amplitude (au) | 1.67 | 1.22 | - | 2.13 | 1.86 | 1.42 | - | 2.3 | 0.19 | 0.5609 |
| Acrophase (ZT) | 1.24 | 1 | - | 1.49 | 1.2 | 0.96 | - | 1.45 | -0.04 | 0.8166 |
| ***SCN mBmal1*** | | | | | | | | | | |
| Mean (au) | 1.89 | 1.82 | - | 1.97 | 1.93 | 1.86 | - | 2.07 | - | - |
| Amplitude (au) | 0.18 | 0.08 | - | 0.29 | 0.23 | 0.12 | - | 0.34 | 0.05 | 0.5112 |
| Phase (ZT) | 12.67 | 12.11 | - | 13.22 | 12.78 | 12.35 | - | 13.21 | 0.12 | 0.7466 |
| ***DG mPer1*** | | | | | | | | | | |
| Mean (au) | 3.78 | 3.46 | - | 4.1 | 3.45 | 2.97 | - | 3.92 | - | - |
| Amplitude (au) | *0.12* | *-0.33* | - | *0.56* | *0.48* | *0* | - | *0.96* | 0.37 | 0.2725 |
| Acrophase (ZT) | *-1.03* | *-5.02* | - | *2.97* | *0.16* | *-0.88* | - | *1.21* | 1.19 | 0.5724 |
| ***BLA mPer1*** | | | | | | | | | | |
| Mean (au) | 2.06 | 1.92 | - | 2.21 | 2.04 | 1.82 | - | 2.25 | - | - |
| Amplitude (au) | *0.05* | *-0.15* | - | *0.25* | 0.23 | 0.01 | - | 0.45 | 0.18 | 0.2347 |
| Phase (ZT) | *13.3* | *8.77* | - | *17.82* | 11.39 | 10.3 | - | 12.47 | -1.91 | 0.4208 |

**Table S4.** Results for the linear models with mixed effects of the percent of diurnal activity in mice lacking expression of *Bmal1* in the forebrain, and their genetic controls, subjected to DF. Statistics correspond to Figs. 3C and Figs S4C.

| Behavior | Genotype | n | Stage | Mean % of diurnal activity | SEM | CI 95% | | | Tukey | LMEM-AOV | |
|---|---|---|---|---|---|---|---|---|---|---|---|
| Feeding | Cami-Bmal +/+ | 13 | Baseline | 24.2 | 2.5 | 18.8 | - | 29.6 | a | | |
| | | | Shocks | 59.2 | 2.9 | 52.9 | - | 65.5 | b | | |
| | | | Post | 55.7 | 2.4 | 50.5 | - | 60.9 | b | | |
| | Cami-Bmal $^{+/-}$ | 9 | Baseline | 26.7 | 2.1 | 21.9 | - | 31.5 | a | Stage: | **2.20E-16** |
| | | | Shocks | 64.9 | 3.3 | 57.1 | - | 72.6 | b | Gen: | **0.0362** |
| | | | Post | 67.4 | 2.6 | 61.3 | - | 73.5 | b | Stage*Gen: | **2.42E-08** |
| | Cami-Bmal $^{-/-}$ | 10 | Baseline | 42.7 | 1.2 | 40 | - | 45.5 | a | | |
| | | | Shocks | 61.5 | 3.5 | 53.7 | - | 69.3 | b | | |
| | | | Post | 53.7 | 1.1 | 51.2 | - | 56.2 | c | | |
| Foraging | Cami-Bmal $^{+/+}$ | 13 | Baseline | 20.5 | 1.7 | 16.9 | - | 24.2 | a | | |
| | | | Shocks | 53.5 | 4.2 | 44.5 | - | 62.6 | b | | |
| | | | Post | 54.6 | 3.4 | 47.3 | - | 61.9 | b | | |
| | Cami-Bmal $^{+/-}$ | 9 | Baseline | 19.6 | 2.2 | 14.5 | - | 24.7 | a | Stage: | **2.00E-16** |
| | | | Shocks | 56.6 | 5.9 | 43 | - | 70.1 | b | Gen: | 0.3236 |
| | | | Post | 67.2 | 3.6 | 58.8 | - | 75.6 | c | Stage*Gen: | **3.15E-02** |
| | Cami-Bmal $^{-/-}$ | 10 | Baseline | 26.6 | 1.3 | 23.5 | - | 29.6 | a | | |
| | | | Shocks | 56.2 | 3.4 | 48.5 | - | 63.9 | b | | |
| | | | Post | 54.7 | 0.5 | 53.6 | - | 55.9 | b | | |
| Nest activity | Cami-Bmal $^{+/+}$ | 13 | Baseline | 28.8 | 1.7 | 25 | - | 32.5 | a | | |
| | | | Shocks | 38.2 | 2.2 | 33.4 | - | 42.9 | b | | |
| | | | Post | 43.7 | 2.1 | 39.2 | - | 48.3 | c | | |
| | Cami-Bmal $^{+/-}$ | 9 | Baseline | 27.9 | 1.4 | 24.7 | - | 31.2 | a | Stage: | **2.00E-16** |
| | | | Shocks | 36.1 | 2.9 | 29.4 | - | 42.8 | b | Gen: | **0.0162** |
| | | | Post | 47.7 | 2.4 | 42.2 | - | 53.2 | c | Stage*Gen: | 0.2123 |
| | Cami-Bmal $^{-/-}$ | 9 | Baseline | 33.5 | 1.3 | 30.6 | - | 36.4 | a | | |
| | | | Shocks | 42 | 1.4 | 38.7 | - | 45.2 | b | | |
| | | | Post | 53.4 | 0.9 | 51.3 | - | 55.5 | c | | |

**Table S5.** Results for the linear models with mixed effects of the amplitude of the fast-Fourier transform (FFT) in mice lacking expression of *Bmal1* in the forebrain, and their genetic controls, subjected to DF. Statistics correspond to Figs. 3D and Figs S4D.

| Behavior | Genotype | n | Stage | Mean FFT amplitude | SEM | CI 95% | | | Tukey | LMEM-AOV | |
|---|---|---|---|---|---|---|---|---|---|---|---|
| Feeding | Cami-Bmal $^{+/+}$ | 13 | Baseline | 0.0096 | 0.0042 | 0.0006 | - | 0.0187 | a | | |
| | | | Shocks | 0.0116 | 0.0024 | 0.0064 | - | 0.0168 | a | | |
| | | | Post | 0.0056 | 0.0031 | -0.0011 | - | 0.0123 | a | | |
| | Cami-Bmal $^{+/-}$ | 9 | Baseline | 0.0112 | 0.0032 | 0.0038 | - | 0.0185 | a | Stage: | **0.0021** |
| | | | Shocks | 0.0182 | 0.0061 | 0.0042 | - | 0.0322 | a | Gen: | 0.3426 |
| | | | Post | 0.0122 | 0.0057 | -0.001 | - | 0.0254 | a | Stage*Gen: | 0.5382 |
| | Cami-Bmal $^{-/-}$ | 10 | Baseline | 0.0044 | 0.002 | -0.0002 | - | 0.009 | a | | |
| | | | Shocks | 0.0144 | 0.005 | 0.0032 | - | 0.0257 | b | | |
| | | | Post | 0.0027 | 0.0009 | 0.0006 | - | 0.0048 | a | | |
| Foraging | Cami-Bmal $^{+/+}$ | 13 | Baseline | 0.034 | 0.0079 | 0.0168 | - | 0.0512 | a | | |
| | | | Shocks | 0.026 | 0.005 | 0.0152 | - | 0.0368 | a | | |
| | | | Post | 0.0158 | 0.004 | 0.0071 | - | 0.0245 | a | | |
| | Cami-Bmal $^{+/-}$ | 9 | Baseline | 0.0697 | 0.0223 | 0.0183 | - | 0.121 | a | Stage: | **0.001** |
| | | | Shocks | 0.0341 | 0.0089 | 0.0135 | - | 0.0546 | b | Gen: | 0.0907 |
| | | | Post | 0.0353 | 0.0169 | -0.0037 | - | 0.0742 | b | Stage*Gen: | 0.2316 |
| | Cami-Bmal $^{-/-}$ | 10 | Baseline | 0.0287 | 0.0101 | 0.0059 | - | 0.0516 | a | | |
| | | | Shocks | 0.0294 | 0.0096 | 0.0077 | - | 0.051 | a | | |
| | | | Post | 0.0053 | 0.0011 | 0.0027 | - | 0.0079 | a | | |
| Nest activity | Cami-Bmal $^{+/+}$ | 13 | Baseline | 0.04 | 0.0088 | 0.0207 | - | 0.0592 | a | | |
| | | | Shocks | 0.0114 | 0.0027 | 0.0055 | - | 0.0173 | b | | |
| | | | Post | 0.0234 | 0.0068 | 0.0086 | - | 0.0381 | a,b | | |
| | Cami-Bmal $^{+/-}$ | 9 | Baseline | 0.0622 | 0.0211 | 0.0134 | - | 0.1109 | a | Stage: | **3.69E-05** |
| | | | Shocks | 0.026 | 0.0105 | 0.0018 | - | 0.0502 | b | Gen: | 0.3173 |
| | | | Post | 0.0444 | 0.0165 | 0.0065 | - | 0.0824 | a,b | Stage*Gen: | 0.1487 |
| | Cami-Bmal $^{-/-}$ | 10 | Baseline | 0.0357 | 0.011 | 0.0108 | - | 0.0607 | a | | |
| | | | Shocks | 0.028 | 0.0077 | 0.0107 | - | 0.0453 | a | | |
| | | | Post | 0.019 | 0.0078 | 0.0013 | - | 0.0367 | a | | |

**Table S6.** Results for the linear models with mixed effects of the percent of activity during the safe phase in mice lacking expression of *Bmal1* in the forebrain, and their genetic controls, subjected to non-cued fear under DD. Statistics correspond to Figs. 3G and Figs S5C.

| Behavior | Genoptype | n | Stage | Mean % of safe-time activity | SEM | CI 95% | | | Tukey | LMEM-AOV | |
|---|---|---|---|---|---|---|---|---|---|---|---|
| Feeding | Cami-Bmal $^{+/+}$ | 8 | Baseline | 26.4 | 1.9 | 22 | - | 30.8 | a | | |
| | | | Shocks | 66.8 | 4.5 | 56.1 | - | 77.5 | b | | |
| | | | Post | 56.6 | 9 | 35.2 | - | 77.9 | b | | |
| | Cami-Bmal $^{+/-}$ | 5 | Baseline | 29.9 | 3.5 | 20.1 | - | 39.8 | a | Stage: | **1.56E-08** |
| | | | Shocks | 70.3 | 0.8 | 68 | - | 72.6 | b | Gen: | 0.8596 |
| | | | Post | 52.5 | 3.1 | 43.8 | - | 61.1 | c | Stage*Gen: | **0.0052** |
| | Cami-Bmal $^{-/-}$ | 5 | Baseline | 48.4 | 0.5 | 46.9 | - | 49.8 | a | | |
| | | | Shocks | 57 | 4.3 | 45.1 | - | 68.9 | a | | |
| | | | Post | 52.7 | 0.7 | 50.9 | - | 54.5 | a | | |
| Foraging | Cami-Bmal $^{+/+}$ | 8 | Baseline | 23.2 | 1.5 | 19.7 | - | 26.6 | a | | |
| | | | Shocks | 71.8 | 3.4 | 63.8 | - | 79.9 | b | | |
| | | | Post | 63.5 | 7.3 | 46.2 | - | 80.8 | b | | |
| | Cami-Bmal $^{+/-}$ | 5 | Baseline | 25 | 2 | 19.5 | - | 30.5 | a | Stage: | **4.82E-14** |
| | | | Shocks | 69.9 | 3.2 | 60.9 | - | 78.8 | b | Gen: | 0.66 |
| | | | Post | 55.9 | 2.8 | 48 | - | 63.7 | c | Stage*Gen: | **8.50E-08** |
| | Cami-Bmal $^{-/-}$ | 7 | Baseline | 46.8 | 0.7 | 45.1 | - | 48.5 | a | | |
| | | | Shocks | 52 | 1.6 | 48 | - | 56 | a | | |
| | | | Post | 51.3 | 0.8 | 49.4 | - | 53.2 | a | | |
| Nest activity | Cami-Bmal $^{+/+}$ | 8 | Baseline | 30.9 | 2.4 | 25.3 | - | 36.5 | a | | |
| | | | Shocks | 43.7 | 3.4 | 35.7 | - | 51.7 | b | | |
| | | | Post | 46.1 | 4.2 | 36.3 | - | 56 | b | | |
| | Cami-Bmal $^{+/-}$ | 5 | Baseline | 26.9 | 1.5 | 22.7 | - | 31 | a | Stage: | **2.10E-11** |
| | | | Shocks | 47.6 | 2.5 | 40.7 | - | 54.5 | b | Gen: | **0.0082** |
| | | | Post | 46.3 | 1.8 | 41.4 | - | 51.2 | b | Stage*Gen: | **0.0006** |
| | Cami-Bmal $^{-/-}$ | 7 | Baseline | 46.3 | 0.5 | 45.1 | - | 47.6 | a | | |
| | | | Shocks | 50.9 | 0.6 | 49.4 | - | 52.4 | a | | |
| | | | Post | 51.4 | 1.1 | 48.7 | - | 54.1 | a | | |

**Table SS7.** Results for the linear models with mixed effects of the amplitude of the fast-Fourier transform (FFT) in mice lacking expression of *Bmal1* in the forebrain, and their genetic controls, subjected to non-cued fear in DD. Statistics correspond to Figs. 3H and Figs S5D.

| Behavior | Genoptype | n | Stage | Mean FFT amplitude | SEM | CI 95% | | | Tukey | LMEM-AOV | |
|---|---|---|---|---|---|---|---|---|---|---|---|
| Feeding | Cami-Bmal $^{+/+}$ | 8 | Baseline | 0.0203 | 0.0077 | 0.0021 | - | 0.0384 | a | | |
| | | | Shocks | 0.0257 | 0.0103 | 0.0014 | - | 0.05 | a | | |
| | | | Post | 0.0254 | 0.0094 | 0.0031 | - | 0.0477 | a | | |
| | Cami-Bmal $^{+/-}$ | 5 | Baseline | 0.0321 | 0.0116 | 0 | - | 0.0642 | a | Stage: | 0.3582 |
| | | | Shocks | 0.0385 | 0.0071 | 0.0189 | - | 0.0581 | a | Gen: | 0.0801 |
| | | | Post | 0.0139 | 0.0081 | -0.0085 | - | 0.0363 | a | Stage*Gen: | 0.3425 |
| | Cami-Bmal $^{-/-}$ | 5 | Baseline | 0.0127 | 0.0092 | -0.0129 | - | 0.0383 | a | | |
| | | | Shocks | 0.0028 | 0.0024 | -0.0037 | - | 0.0093 | a | | |
| | | | Post | 0.0028 | 0.0011 | -0.0003 | - | 0.0059 | a | | |
| Foraging | Cami-Bmal $^{+/+}$ | 8 | Baseline | 0.0639 | 0.0124 | 0.0345 | - | 0.0932 | a | | |
| | | | Shocks | 0.0906 | 0.0205 | 0.0423 | - | 0.139 | a | | |
| | | | Post | 0.0712 | 0.019 | 0.0263 | - | 0.1162 | a | | |
| | Cami-Bmal $^{+/-}$ | 5 | Baseline | 0.0778 | 0.0204 | 0.0212 | - | 0.1344 | a | Stage: | 0.0996 |
| | | | Shocks | 0.0782 | 0.0085 | 0.0547 | - | 0.1017 | a | Gen: | **0.0003** |
| | | | Post | 0.0325 | 0.009 | 0.0074 | - | 0.0576 | b | Stage*Gen: | 0.1088 |
| | Cami-Bmal $^{-/-}$ | 7 | Baseline | 0.0066 | 0.002 | 0.0017 | - | 0.0116 | a | | |
| | | | Shocks | 0.0039 | 0.0009 | 0.0018 | - | 0.006 | a | | |
| | | | Post | 0.0076 | 0.0025 | 0.0016 | - | 0.0136 | a | | |
| Nest activity | Cami-Bmal $^{+/+}$ | 8 | Baseline | 0.0675 | 0.0157 | 0.0304 | - | 0.1045 | a | | |
| | | | Shocks | 0.025 | 0.0039 | 0.0158 | - | 0.0342 | b | | |
| | | | Post | 0.0356 | 0.0102 | 0.0114 | - | 0.0598 | a,b | | |
| | Cami-Bmal $^{+/-}$ | 5 | Baseline | 0.0985 | 0.0136 | 0.0607 | - | 0.1363 | a | Stage: | **0.0004** |
| | | | Shocks | 0.0311 | 0.0082 | 0.0083 | - | 0.0539 | b | Gen: | **3.84E-06** |
| | | | Post | 0.0601 | 0.0264 | -0.0131 | - | 0.1333 | a,b | Stage*Gen: | 0.113 |
| | Cami-Bmal $^{-/-}$ | 7 | Baseline | 0.0128 | 0.0032 | 0.0051 | - | 0.0206 | a | | |
| | | | Shocks | 0.007 | 0.002 | 0.002 | - | 0.012 | a | | |
| | | | Post | 0.0126 | 0.0031 | 0.0051 | - | 0.0201 | a | | |

**Table S8.** Results for the linear models with mixed effects of the amplitude of the fast-Fourier transform (FFT) in mice with targeted deletion of *Bmal1* to the SCN, and their controls, subjected to non-cued fear in DD. Statistics correspond to Figs. 4C and Figs S6D.

| Behavior | Genotype | n | Stage | Mean FFT amplitude | SEM | CI 95% | Tukey | LMM-AOV | |
|---|---|---|---|---|---|---|---|---|---|
| Feeding | SCN-Bmal1 $^{-/-}$ | 6 | Baseline | 0.0043 | 0.0005 | 0.0030 - 0.0055 | a | Stage: | 0.100676 |
| | | | Shocks | 0.0062 | 0.0009 | 0.0039 - 0.0086 | a | | |
| | | | Post | 0.0160 | 0.0049 | 0.0034 - 0.0286 | a | Gen: | **0.039836** |
| | SCN-Bmal1 $^{+/+}$ | 4 | Baseline | 0.0308 | 0.0122 | -0.0081 - 0.0697 | a,b | | |
| | | | Shocks | 0.0427 | 0.0145 | -0.0035 - 0.0890 | a | Stage*Gen: | **0.000897** |
| | | | Post | 0.0148 | 0.0049 | -0.0009 - 0.0305 | b | | |
| Foraging | SCN-Bmal1 $^{-/-}$ | 6 | Baseline | 0.0068 | 0.0026 | 0.0001 - 0.0135 | a | Stage: | 0.053748 |
| | | | Shocks | 0.0112 | 0.0049 | -0.0014 - 0.0238 | a | | |
| | | | Post | 0.0154 | 0.0071 | -0.0027 - 0.0336 | a | Gen: | **0.005911** |
| | SCN-Bmal1 $^{+/+}$ | 4 | Baseline | 0.0510 | 0.0127 | 0.0104 - 0.0916 | a | | |
| | | | Shocks | 0.0595 | 0.0151 | 0.0115 - 0.1075 | a | Stage*Gen: | **0.007886** |
| | | | Post | 0.0221 | 0.0064 | 0.0017 - 0.0425 | b | | |
| Nest activity | SCN-Bmal1 $^{-/-}$ | 6 | Baseline | 0.0115 | 0.0033 | 0.0029 - 0.0201 | a | Stage: | 0.05212 |
| | | | Shocks | 0.0066 | 0.0018 | 0.0019 - 0.0112 | a | | |
| | | | Post | 0.0063 | 0.0024 | 0.0001 - 0.0125 | a | Gen: | 0.10213 |
| | SCN-Bmal1 $^{+/+}$ | 4 | Baseline | 0.0306 | 0.0157 | -0.0194 - 0.0807 | a | | |
| | | | Shocks | 0.0178 | 0.0062 | -0.0021 - 0.0376 | a,b | Stage*Gen: | 0.27096 |
| | | | Post | 0.0073 | 0.0018 | 0.0015 - 0.0131 | b | | |

**Table S9.** Results for Raleigh statistics on the phase of mice with restricted expression of *Bmal1* to the SCN and subjected to non-cued fear under DD. Statistics correspond to Fig. 4F and S7D.

| | | | Rayleigh statistics | | | | |
|---|---|---|---|---|---|---|---|
| Behavior | Stage | n | r | p | Phase (h) | Fiduciary Limits | SEM |
| Feeding | Post/ Baseline | 11 | 0.73 | 0.0013 | 0.80 | 21.68 - 0.72 | 0.84 |
| | Post/Shocks | 11 | 0.48 | 0.079 | 1.02 | 25.11 - 3.16 | 1.18 |
| Foraging | Post/ Baseline | 11 | 0.69 | 0.0029 | 0.60 | 22.98 - 2.23 | 0.89 |
| | Post/Shocks | 11 | 0.03 | 0.988 | 18.3 | 15.4 - 21.2 | 1.60 |
| Nest activity | Post/ Baseline | 11 | 0.68 | 0.0036 | 0.15 | 22.2 - 1.51 | 0.91 |
| | Post/Shocks | 11 | 0.24 | 0.526 | 10.29 | 7.73 - 12.86 | 1.41 |
